# The plant longevity gene *AHL15* delays leaf senescence by repressing *ORESARA1* and cytokinin degradation

**DOI:** 10.1101/2025.11.23.689984

**Authors:** Thalia Luden, Petra Amakorová, Ondřej Novák, Salma Balazadeh, Remko Offringa

## Abstract

The *Arabidopsis thaliana* (Arabidopsis) *AT-HOOK MOTIF NUCLEAR LOCALIZED 15 (AHL15)* gene is associated with various longevity phenotypes and extends the life span of plants when overexpressed. In this study, we show that, in addition to previously described longevity phenotypes, constitutive overexpression of *AHL15* in Arabidopsis delays leaf senescence, whereas *ahl15* loss-of-function accelerates this process. Dexamethasone-induced nuclear localization of AHL15-GR during dark-triggered senescence results in a stay-green phenotype and represses the expression of several early senescence-associated genes. Among these, the *ORESARA1 (ORE1)* locus is directly bound by AHL15, suggesting a direct repressive effect of AHL15 on senescence. Furthermore, we demonstrate that AHL15 acts by directly repressing the expression of several *CYTOKININ OXIDASE (CKX)* genes involved in cytokinin inactivation, resulting in a delayed degradation of cytokinins during dark-induced senescence. Cytokinins are known to delay senescence, and together with the downregulation of *ORE1* expression, this explains the repressive effect of AHL15 on senescence.

**Significance:** Here, we show that in addition to its previously reported effects on aging processes, the *AT-HOOK MOTIF NUCLEAR LOCALIZED*-family protein AHL15 also represses leaf senescence. Our results demonstrate that AHL15 delays the senescence program in two ways: by directly repressing the senescence master regulator *ORESARA1,* and by transcriptional repression of *CYTOKININ OXIDASE* genes, resulting in a delayed breakdown of the senescence-inhibiting hormone cytokinin, which together explain the strong stay-green phenotype of *AHL15*-overexpressing plants.

## Introduction

Leaf senescence is the process during which the photosynthetic machinery of leaves is catabolized and recycled for use in other parts of the plant (Lim, Kim and Nam, 2007; Schippers *et al*., 2015; Guo *et al*., 2021). It is the final developmental stage of leaves, and its progression is tightly controlled by various transcription factors (TFs). Based on research primarily conducted in the model plant *Arabidopsis thaliana* (Arabidopsis), the TFs ORESARA1 (ORE1) and NAC-LIKE, ACTIVATED BY AP3/PI (AtNAP) have been identified as major regulators of senescence (Guo and Gan, 2006; Kim *et al*., 2009; Qiu *et al*., 2015). Loss-of-function of *ore1*, *nap,* or other positive regulators of senescence results in a stay-green phenotype (Guo and Gan, 2006; Kim *et al*., 2009), and ORE1 regulates the transcription of numerous genes in the senescence pathway (Balazadeh *et al*., 2010). Upon initiation of senescence, genes involved in chlorophyll degradation, also referred to as chlorophyll catabolic genes (CGGs) are upregulated by ORE1, NAP, and other TFs in the senescence pathway, and together the resulting enzymes degrade chlorophyll (Chl) into smaller molecules that can be exported to other tissues in the plant, resulting in the characteristic yellowing of senescent leaves (Qiu *et al*., 2015; Yang, Worley and Udvardi, 2015; Woo *et al*., 2019). The remobilization of nutrients from older leaves to other parts of the plant helps regulate nutrient availability and distribution, thereby increasing the plant’s chances of survival during periods of nutrient scarcity (Schippers *et al*., 2015). However, when senescence negatively impacts the overall quality of a crop, such as in leafy vegetables or in ornamental plants, delayed senescence can be a desirable trait. Therefore, improving the control of leaf senescence is essential for enhancing crop quality.

Senescence can be induced by various factors, including leaf age, dark incubation, nutrient shortage, and the application of hormones such as ethylene, jasmonic acid, abscisic acid, salicylic acid, or strigolactones (Woo *et al*., 2019). In Arabidopsis, ethylene triggers a signaling cascade that activates the transcription factor ETHYLENE INSENSITIVE 3 (EIN3), which in turn promotes the transcription of *ORE1* and *NAP* (Li *et al*., 2013; Kim *et al*., 2014). Additionally, dark incubation induces transcription of both *ORE1* and *EIN3* via PHYTOCHROME INTERACTING FACTOR4 (PIF4) and PIF5 (Song *et al*., 2014). In monocarpic plants, flowering also triggers leaf senescence, allowing the relocation of nutrients from leaves to seeds and thereby enhancing the offspring’s chances of survival (Thomas, 2013). For example, in Arabidopsis, the whole plant senesces within a few weeks after flowering. This process is controlled by several TFs, among which MYB2 and FRUITFULL (FUL), and coincides with reduced production of the hormone cytokinin (CK) (Guo and Gan, 2011; Balanzà *et al*., 2018).

In contrast to most other plant hormones that induce senescence, CKs repress leaf senescence: exogenous CK application significantly delays leaf senescence in various plant species (Richmond *et al*., 1957; Gan and Amasino, 1995; Hallmark and Rashotte, 2020; Pokorná *et al*., 2021). The senescence-delaying activity of CK is mediated by the CK receptor ARABIDOPSIS HISTIDINE KINASE 3 (AHK3), which phosphorylates ARABIDOPSIS RESPONSE REGULATOR 2 (ARR2) upon CK perception (Kim *et al*., 2006). ARR2 then promotes expression of *CYTOKININ RESPONSE FACTOR 6* (*CRF6*), a TF that negatively regulates leaf senescence via an unknown mechanism (Zwack *et al*., 2013). Loss-of-function mutants of *ahk3* show accelerated senescence phenotypes and the senescence-delaying effect of CK treatment is reduced in these plants as a result of their weakened ability to transduce CK signals (Kim *et al*., 2006). A similar phenotype can be seen in *crf6* mutants (Zwack *et al*., 2013).

CKs are synthesized predominantly by enzymes of the ISOPENTENYL TRANSFERASE (IPT) and LONELY GUY (LOG) families. IPT-family enzymes are responsible for adding a prenyl group to N^6^ of ATP or ADP (Kakimoto, 2001; Takei, Sakakibara and Sugiyama, 2001). Additional enzymes such as CYP735A are involved in making changes to this prenyl group, thereby producing precursors to different CK variants (Argueso and Kieber, 2024). Biological activation of these CK precursors relies on the removal of the riboside group by LOG-family enzymes, which are highly redundant and of which LOG7 and LOG8 appear to be the most important in Arabidopsis (Kuroha *et al*., 2009; Tokunaga *et al*., 2012). CK degradation is predominantly accomplished by enzymes of the CYTOKININ OXIDASE/DEHYDROGENASE (CKX) class. CKX enzymes irreversibly cleave off the N^6^ side chains from active CKs, thereby inactivating them permanently (Galuszka *et al*., 2000; Schmülling *et al*., 2003). Knockout of the rice *ckx11* gene causes a delay in leaf senescence, indicating that reduced CK degradation negatively affects senescence (W. Zhang *et al*., 2021). On the other hand, overexpression of *LOG4* delays leaf senescence in Arabidopsis (Kuroha et al., 2009) and causes pleiotropic effects. In addition, ectopic expression of the *Agrobacterium tumefaciens IPT* gene under the senescence-specific *SAG12* promoter delays leaf senescence in several crops (Gan and Amasino, 1995; Jordi *et al*., 2000; McCabe *et al*., 2001). Thus, leaf senescence can be regulated by controlling the expression of the CK biosynthesis genes of the *IPT* and *LOG* families and by regulating the expression of the CK-degrading *CKX* genes.

While CK plays an important role in senescence, the regulatory mechanisms that control CK-mediated inhibition of senescence remain elusive. So far, only two TFs that regulate CK levels during senescence have been identified: MYB2 and NAP (Guo and Gan, 2011; Hu *et al*., 2021; Li *et al*., 2023). In wild type plants, *IPT* expression in the leaf is silenced by MYB2, whose expression increases with progressing age (Guo and Gan, 2011). It was shown that *MYB2* expression in Arabidopsis and the legume *Caragana intermedia* can be repressed by proteins of the S40 family, and overexpression of *S40*-family genes results in enhanced CK levels and a stay-green phenotype (Yang *et al*., 2022). The expression of *CRF6* also decreases in older leaves through an unknown mechanism (Zwack *et al*., 2013). At the same time, *CKX3* expression is induced by the senescence-promoting TF NAP, resulting in enhanced CK degradation (Hu *et al*., 2021). The repression of *CKX* expression by NAP was observed in Arabidopsis and Chinese cabbage, indicating that this pathway is conserved (Hu *et al*., 2021; Li *et al*., 2023). These results show a clear link between CK and age-induced senescence, but additional regulatory mechanisms that control the expression of CK biosynthesis- and degrading genes during the senescence process remain to be identified.

Previously, we identified the Arabidopsis gene *AT-HOOK MOTIF NUCLEAR LOCALIZED15 (AHL15)* as a regulator of plant longevity: overexpression of *AHL15* in Arabidopsis resulted in an increased life span and flowering time and enhanced vegetative development of axillary meristems, branching and secondary growth, whereas *ahl15* loss-of-function or perturbed *ahl* function reduced all these aspects (Karami *et al*., 2020; Rahimi, Karami, Lestari, *et al*., 2022). CK levels were increased in stems of *AHL15* overexpressing plants, explaining the enhanced secondary growth and branching phenotypes in these plants (Rahimi, Karami, Lestari, *et al*., 2022). In Arabidopsis, the *AHL* gene family is comprised of 29 members that are subdivided into two clades (clades A and B) (Zhao *et al*., 2014). Previous research has demonstrated that members of the clade A *AHL* subfamily, including *AHL15*, exhibit a high level of functional redundancy (Yun *et al*., 2012; Zhao *et al*., 2013; Karami *et al*., 2020). Lim et al. (2007) showed that overexpression of the clade A member *AHL27* delays leaf senescence, whereas a recent study has identified the clade B member *AHL9* as a promoter of leaf senescence, suggesting that clade A and B AHLs may function antagonistically (Lim *et al*., 2007; Zhou *et al*., 2022).

Here, we show that, similar to *AHL27,* overexpression of *AHL15* delays leaf senescence, while loss-of-function *ahl15* mutants exhibit early senescence. Our results show that the expression of several senescence marker genes, including *ORESARA1* (*ORE1*), is repressed when AHL15 activity is induced and that AHL15 directly binds to the *ORE1* locus, indicating that AHL15 directly suppresses the senescence program. Additionally, we found that AHL15 binds upstream of the *CKX2*, *CKX3*, and *CKX5* genes and that induction of AHL15 activity leads to nearly complete silencing of their expression. This silencing coincides with a slower decline in CK levels during dark-induced senescence. Together, our data reveal that AHL15 delays the senescence program through parallel pathways: by repressing the expression of *ORE1* and by repressing *CKX* expression and subsequent CK degradation.

## Results

### *AHL15* overexpression delays leaf senescence

Given that overexpression of *AHL15* extends overall plant longevity and because delayed senescence has previously been reported for *AHL27* overexpressing lines (Lim *et al*., 2007), we tested whether a similar senescence phenotype could also be observed in *p35S:3xFLAG-AHL15* plants (hereafter referred to as *AHL15* OX). To this end, we compared senescence in the fifth leaf of Col-0 (wild type) and *ahl15* knockout plants with two independent *AHL15* OX lines in both dark-induced (Figure 1) and age-dependent conditions (Figure S1). To quantify senescence, we measured leaf Chlorophyll (Chl) using two methods: 1) Chl extraction using acetone followed by spectrophotometry (Arnon, 1949), and 2) a recently developed colorimetric assay. This assay measures the pixel intensity value in Red, Green, and Blue channels of digital images, which can be used to calculate the normalized Red value (Rn; Red/(Red+Green+Blue)). Previously, we have demonstrated that the Rn value shows a significant negative correlation with the total Chl content of leaves, which makes it an appropriate non-invasive method for Chl quantification (Luden *et al*., 2025).

**Figure 1.**
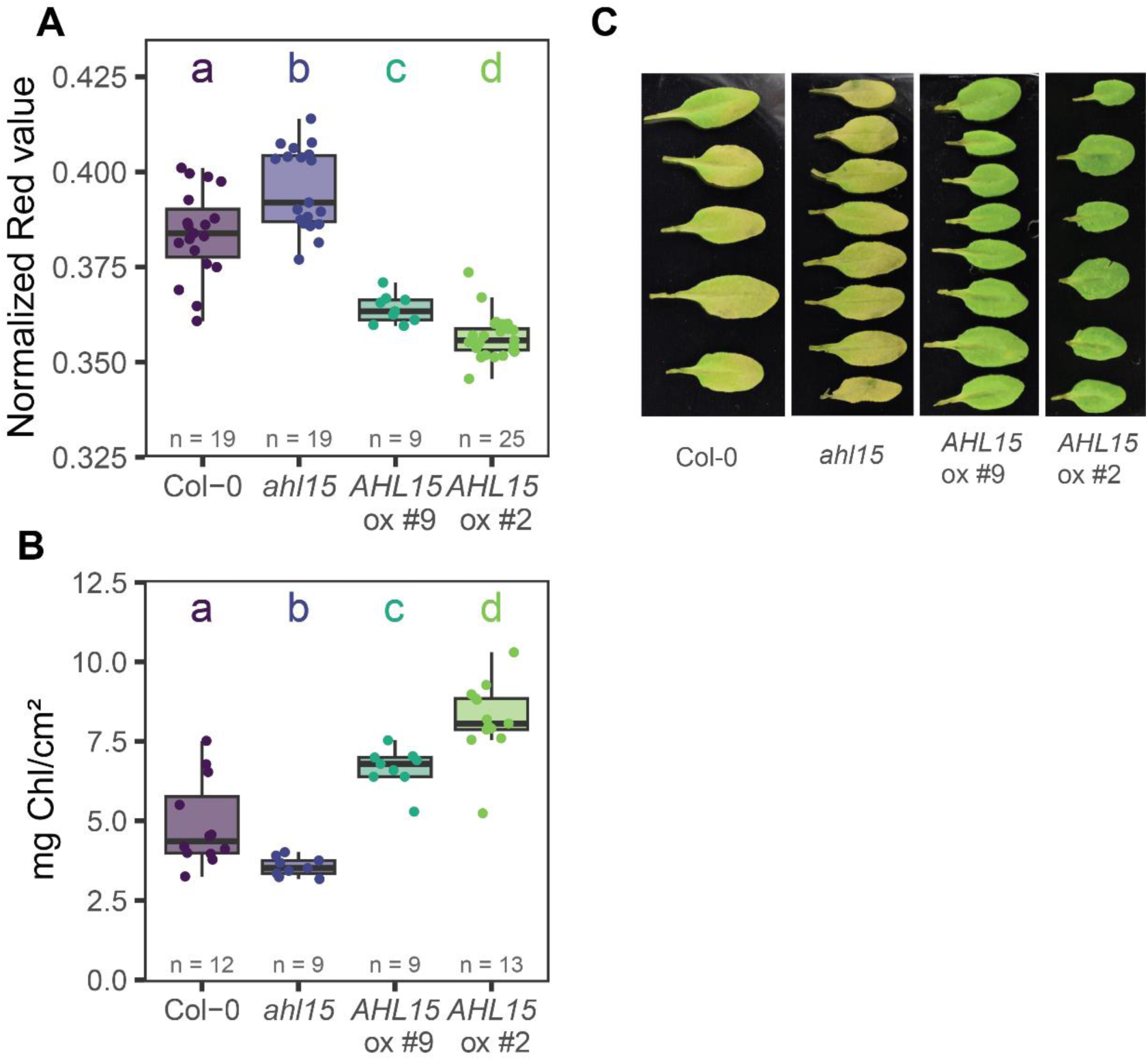
Dark-induced leaf *s*enescence is delayed by *AHL15* overexpression and accelerated by *ahl15* loss-of-function. **A-C:** The fifth leaf of 23-day-old wild-type (Col-0), *ahl15* mutant or *p35S:3xFLAG-AHL15* (*AHL15 OX*) Arabidopsis plants was detached and floated on 5 ml (5 mM) MES buffer for five days in the dark to induce the senescence process. Leaves were imaged for chlorophyll quantification as the normalized Red value (Rn) (**A**). Chlorophyll was extracted and quantified by spectrophotometer (**B**). Representative images of leaves of each genotype (**C**). Differences between genotypes in **A** and **B** were compared by a Kruskal-Wallis test, and significance groups are indicated by letters (n=9-25; p < 0.05). White bars in **C** represent 1 cm.

In dark-induced conditions, the senescence of the fifth leaf was significantly delayed in the *AHL15* OX lines compared to the wild type, as evidenced by both the lower Rn value (Figure 1A) and the higher total Chl content (Figure 1B). In addition, *ahl15* loss-of-function plants showed a mild but significant increase in Rn value and a decrease in total Chl content compared to Col-0. These findings demonstrate that *AHL15* inhibits leaf senescence in dark-induced conditions.

Similarly, at 42 days after germination, the older leaves of wild type and *ahl15* knockout plants started to senesce, whereas leaves of the same age in *AHL15* OX lines stayed green, indicating that age-dependent senescence could be delayed by *AHL15* overexpression (Figure S1A). To monitor this in more detail, we harvested the fifth leaves of different plants at 21, 24, 27, 30, 34, and 36 days of emergence and used images of these leaves to measure the Rn value (Figure S1B, C). This showed that the progression of senescence in the fifth leaf was slightly delayed in *AHL15* OX plants (most strongly in *AHL15* OX line 2) compared to wild type and *ahl15* loss-of-function plants, suggesting that AHL15 may play a role in age-dependent senescence as well. The latter is in line with the expression of *AHL15*, which is on in young developing leaves and turned off when leaves are fully grown (Figure S2)

Overexpression of *AHL15* and other clade-A *AHLs* generally delays flowering time (Street et al., 2008; Xiao et al., 2009; Karami et al., 2020), and AHL15 has been shown to delay vegetative phase change (Rahimi, Karami, Balazadeh, *et al*., 2022). In addition, *AHL15* is expressed primarily in juvenile, newly developing leaves and is absent in older and adult-stage leaves (Rahimi, Karami, Balazadeh, *et al*., 2022). To evaluate the effects of *AHL15* overexpression on the senescence phenotype independently of other developmental processes, we used a dexamethasone (DEX)-inducible *p35S:AHL15-GR* line in which the AHL15-GR fusion protein translocates to the nucleus upon DEX treatment (Karami *et al*., 2020). During the dark incubation period, leaves were floated on 5 mL MES buffer, to which DEX was added to a final concentration of 10 μM to induce AHL15 activity simultaneously with the senescence process. DEX-treated *p35S:AHL15-GR* leaves showed a similar delayed senescence phenotype as *AHL15* OX leaves, whereas mock-treated *p35S:AHL15-GR* leaves senesced like wild-type leaves (Figure 2A-C).

**Figure 2.**
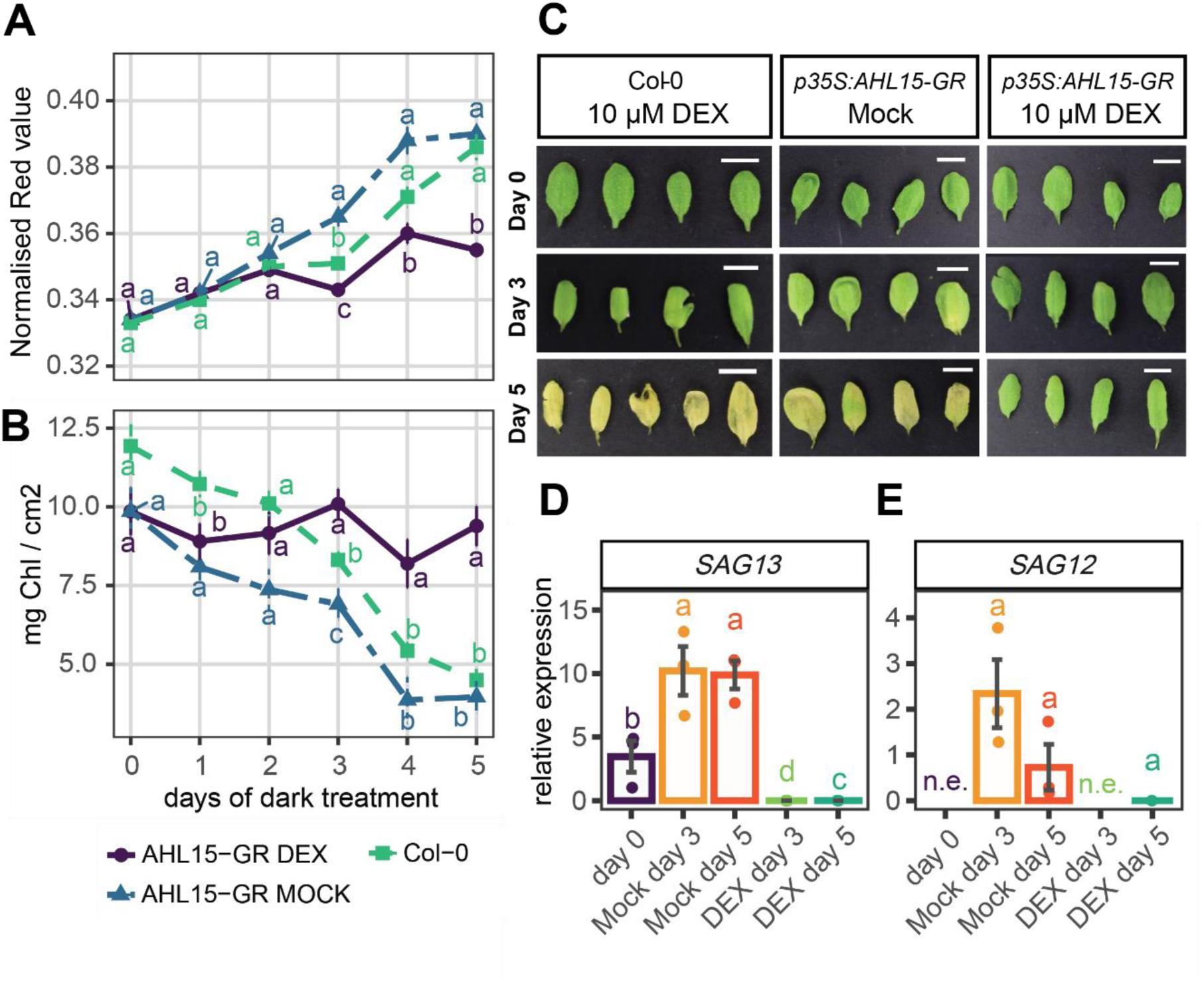
DEX-induced nuclear localization of AHL15-GR inhibits leaf senescence. **A-C:** The fifth leaf was detached from 23-day-old wild-type (Col-0) or *p35S:AHL15-GR* plants and incubated in the dark on 5 mM MES buffer with or without 10 µM DEX for the indicated number of days. The leaf chlorophyll content was quantified over five days of dark incubation by image-based calculation of the normalized Red value (**A**) or by direct chlorophyll extraction (**B**) (n = 8 for days 1, 2, and 4, and n = 16 for days 0, 3, and 5). The quantification was repeated twice with similar results. Representative images of leaves from each group before and after three or five days of dark incubation (**C**). White bars in (**C**) represent 1 cm. **D, E:** Expression of the senescence-associated genes *SAG13* (**D**) and *SAG12* (**E**) in the DEX-or Mock-treated *p35S:AHL15-GR* leaves shown in **C**, as determined by RT-qPCR. RNA was extracted from three samples per time point and condition (n=3), with each sample consisting of the fifth leaf of three individual plants. Expression was normalized to *ACTIN2*. *SAG12* expression could not be detected (n.e. for not expressed) at day 0 and in DEX-treated leaves at day 3. Letters indicate significance groups calculated with the Kruskall-Wallis test (p < 0.05).

We next investigated the effect of DEX-induced AHL15-GR nuclear localization on the expression of senescence-associated genes after increasing periods of dark incubation. Based on the Rn value and total Chl content of leaves incubated in the dark over five days, significant differences between DEX-induced and non-induced leaves (DEX-treated *vs.* mock-treated *p35S:AHL15-GR,* or DEX-treated Col-0) became evident after three days of dark incubation. This indicates that senescence had been initiated at this time point in non-induced leaves (Figure 2A-C). To determine whether AHL15 inhibits senescence at its onset or a later stage, we measured the expression of *SENESCENCE-ASSOCIATED GENES (SAGs) SAG13* and *SAG12,* which are associated with early- and late senescence, respectively (Lohman *et al*., 1994). RNA was isolated from leaves right before dark incubation and DEX treatment (day 0), and after three or five days of dark incubation with and without DEX treatment. A clear and significant difference in the expression of the early senescence marker *SAG13* as well as the late senescence marker *SAG12* was visible between DEX-and mock-treated plants (Figure 2D-E), whereas the expression of *SEN1* was slightly but significantly reduced in DEX-treated leaves after 5 days (Figure S3). As the expression of *SEN1* is induced by darkness (Oh *et al*., 1996; Chung *et al*., 1997), these results indicate that overexpression of *AHL15* predominantly affects the onset of senescence at an early stage and interferes with the response to darkness less strongly.

### AHL15 directly represses the expression of *ORE1* and several *CKX* genes

To understand how AHL15 suppresses leaf senescence, we investigated whether AHL15 binds any senescence genes directly by consulting AHL15 ChIP-seq data from *AHL15* overexpressing plants *(p35S:3xFLAG-AHL15)* that we generated in a different study (Luden, Chouaref and Offringa, 2025). ChIP-seq peaks of AHL15 were present up- and downstream of many genes, among which the *SQUAMOSA PROMOTER BINDING PROTEIN-LIKE 9 (SPL9)* gene (Figure S4), whose expression was previously shown to be suppressed by AHL15 (Rahimi, Karami, Balazadeh, *et al*., 2022). Interestingly, significant peaks were present at 600 bp upstream and 1 kb downstream of the senescence regulator *ORE1,* which could explain the delayed senescence phenotype observed in *AHL15* overexpressing plants (Figure 3A). We next measured the expression of *ORE1* after senescence induction of DEX- or mock-treated leaves and saw that its expression was strongly repressed in DEX-treated leaves, implying a direct repression of this gene by AHL15 (Figure 3B).

**Figure 3.**
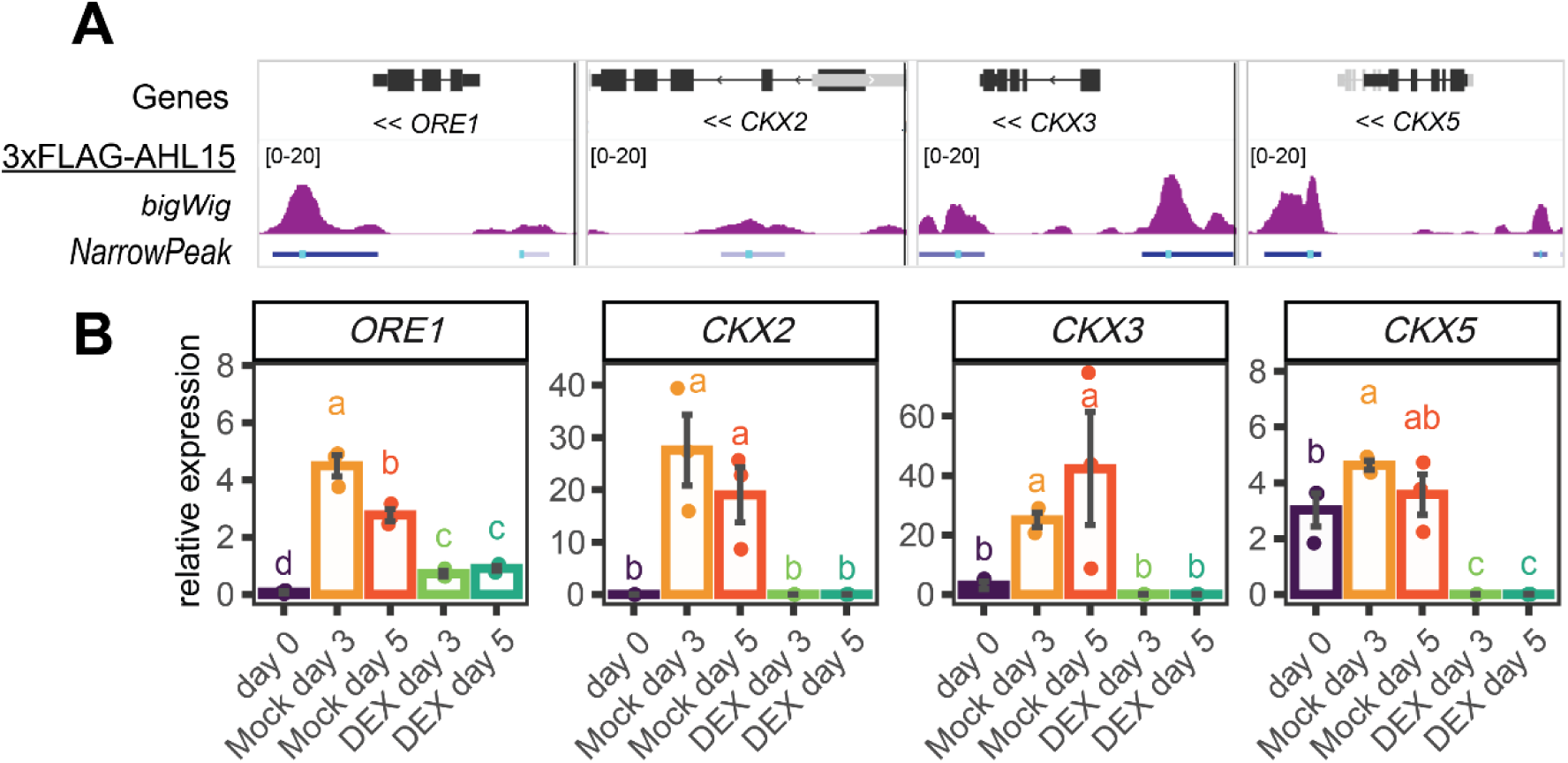
AHL15 binds to the *ORE1* and *CKX* loci to repress gene expression. **A:** ChIP performed on the whole rosette of 31-day-old *p35S:3xFLAG-AHL15* plants using an anti-FLAG antibody. ChIP seq detected peaks of AHL15 binding 1 kb downstream and 600bp upstream of *ORE1*, in the second intron of *CKX2*, 1.9 kb upstream and directly downstream of *CKX3*, and 2.3 kb upstream and 2 kb downstream of *CKX5*. Genes are shown in black (black blocks are exons, black lines are introns, grey blocks represent genes on the opposite strand), read alignments are shown in purple (bigwig track), and the location of significant peaks are indicated in blue, with a light blue bar indicating the peak summit (narrowpeak track). Similar results were obtained in three biological replicates. **B:** The expression of *ORE1*, *CKX2, CKX3* and *CKX5* is repressed by DEX induced AHL15-GR activation in dark-incubated leaves. Leaves of 23-day-old *p35S:AHL15-GR* plants were floated on 5 mM MES buffer with or without 10 µM DEX for the indicated number of days in the dark. RNA used for RT-qPCR was extracted from three samples per time point and condition (n=3), with each sample consisting of the fifth leaf of three individual plants. Expression was normalized to *ACTIN2*. Letters indicate significance groups as calculated by the Kruskall-Wallis test (p < 0.05).

In addition, we identified the *CKX* genes *2*, *3*, and *5* (Figure 3A) and several CK biosynthesis genes (*IPT7, LOG1, LOG7, LOG8;* Figure S5A) among the significant ChIP-seq peaks. Because of the previous finding that AHL15 promotes wood formation by enhancing CK levels (Rahimi, Karami, Lestari, *et al*., 2022), and because CKs are known to have an delaying effect on leaf senescence (Kim *et al*., 2006; Guo and Gan, 2011; Hu *et al*., 2021; Yang *et al*., 2022), we hypothesized that the delayed senescence phenotype of *AHL15* overexpressing plants could in part be a result of increased CK levels. To test whether AHL15 directly regulates the expression of these genes, we measured the expression of each of these genes at 0, 3, or 5 days after senescence induction in DEX- or mock-treated *p35S:AHL15-GR* leaves. The expression of *LOG8* and the three *CKX* genes in mock-treated leaves followed a similar pattern as the senescence markers *SAG13* and *SAG12*, and were all increased after 3 and 5 days of dark incubation (Figure 3B, Figure S5B). In contrast, the expression of *IPT7* and *LOG1* in mock-treated leaves was lower than at day 0, and the expression of *LOG7* was not affected (Figure S5B). Interestingly, the expression of the *CKX* genes was significantly lower in DEX-treated leaves compared to the mock treatment, implying that these genes are directly repressed by AHL15 (Figure 3B). The repression of *IPT7* expression upon dark incubation was delayed in DEX-treated leaves, whereas the expression of *LOG1* and *LOG8* was repressed in comparison to mock-treated leaves and *LOG7* expression was unaffected (Figure S5B). These data suggest that unlike in stem tissue, where AHL15 promotes CK biosynthesis (Rahimi, Karami, Lestari, *et al*., 2022), it delays leaf senescence rather by repressing the expression of cytokinin-degrading CKX proteins.

### Cytokinin levels are increased in *AHL15* overexpressing plants

The repression of *CKX* genes by AHL15 suggests that the inactivation of CKs is delayed in *AHL15-*overexpressing leaves, resulting in increased CK levels that repress the onset of senescence. To test whether this is the case, we measured the levels of a wide range of CKs in dark-induced *AHL15-GR* leaves with and without DEX treatment before and during senescence (Figure 4, Figures S6-S11). These measurements showed that total CK levels drop upon senescence induction in both DEX- and mock-treated leaves, but that this decrease in CK levels is less pronounced in DEX-treated leaves (Figure 4A). This trend was especially clear for the isopentenyladenine (iP)-glucosides (Figure 4B, Figure S6), implying that they may be especially sensitive to degradation by CKX enzymes during the senescence process. Previous research has shown that both iP and iP-9-glucoside (IP9G), but not iP-7-glucoside (iP7G), can inhibit senescence in Arabidopsis cotyledons (Hallmark and Rashotte, 2020). Although iP in its base form was not present at detectable levels in any of the samples, the elevated levels of iP9G in DEX-treated samples may partially explain the delayed senescence phenotype. In addition, the levels of trans-zeatin base and trans-zeatin Riboside (tZ and tZR) were reduced in all dark-incubated samples, but their concentration remained significantly higher in DEX-treated leaves (Figure 4C-D, Figure S7). Of all CK types, tZ-base and tZR are among the most potent inhibitors of senescence (Hönig *et al*., 2018; Pokorná *et al*., 2021), and their elevated concentration in DEX-versus mock-treated leaves could therefore play an important role in senescence inhibition in DEX-treated leaves.

**Figure 4.**
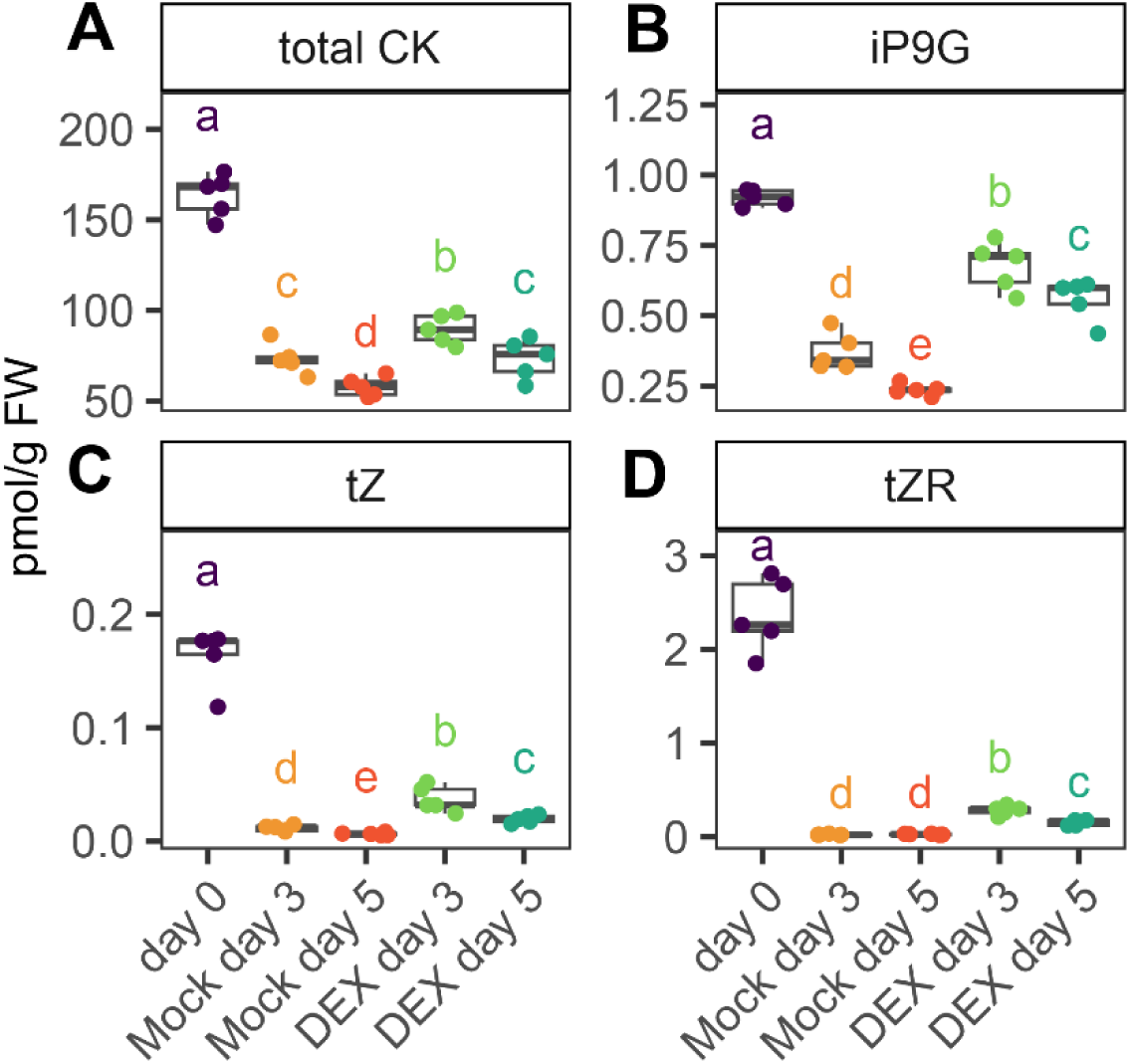
DEX-induced AHL15-GR activation delays the reduction in cytokinin (CK) levels in dark-incubated leaves. **A-D:** Quantification of the total CK (**A**), isopentenyladenine-9-glucoside (**B**), trans-zeatin base (**C**) or trans-zeatin riboside (**D**) level in the fifth leaf of 23-day old *p35S:AHL15-GR* plants floated on 5 mM MES with or without 10 µM DEX in the dark for the indicated number of days. For each sample three leaves were pooled and five samples were measured for each condition (n = 5). Groups were compared using the Kruskall-Wallis test, and letters indicate significance groups (p < 0.05).

In addition to tZ and iP, we profiled the levels of cis-zeatine (cZ; Figure S8) and dihydrozeatin (DHZ; Figure S9) during the senescence process. Interestingly, cZ-base and cZG levels increased upon dark induction and were especially high after five days of DEX treatment (Figure S8). cZ has a weak effect on leaf senescence, and its function in the plant remains largely unknown (Gajdošová *et al*., 2011). Previous research has shown that DHZ levels drop during the senescence process in tobacco leaves (Hönig *et al*., 2018), and we observed a similar trend in our experiments (Figure S9). While DHZ-base levels were undetectable, we found that DHZ derivatives were present at reduced levels in all samples and did not differ significantly between treatments, indicating that this form of CK is not affected by AHL15 or its downstream targets (Figure S9). Finally, we observed an overall increase in CK bases in DEX-treated samples compared to untreated and mock-treated samples (Figure S10), suggesting that other CK bases may also contribute to the AHL15-mediated delay of senescence.

Taken together, our results indicate that AHL15 delays senescence through multiple mechanisms: by directly repressing *ORE1* expression, and by downregulating *CKX* genes, thereby delaying the inactivation of CKs in leaves. The latter process results in elevated CK levels in *AHL15*-overexpressing leaves, ultimately contributing to the delay in leaf senescence (Figure 5).

**Figure 5.**
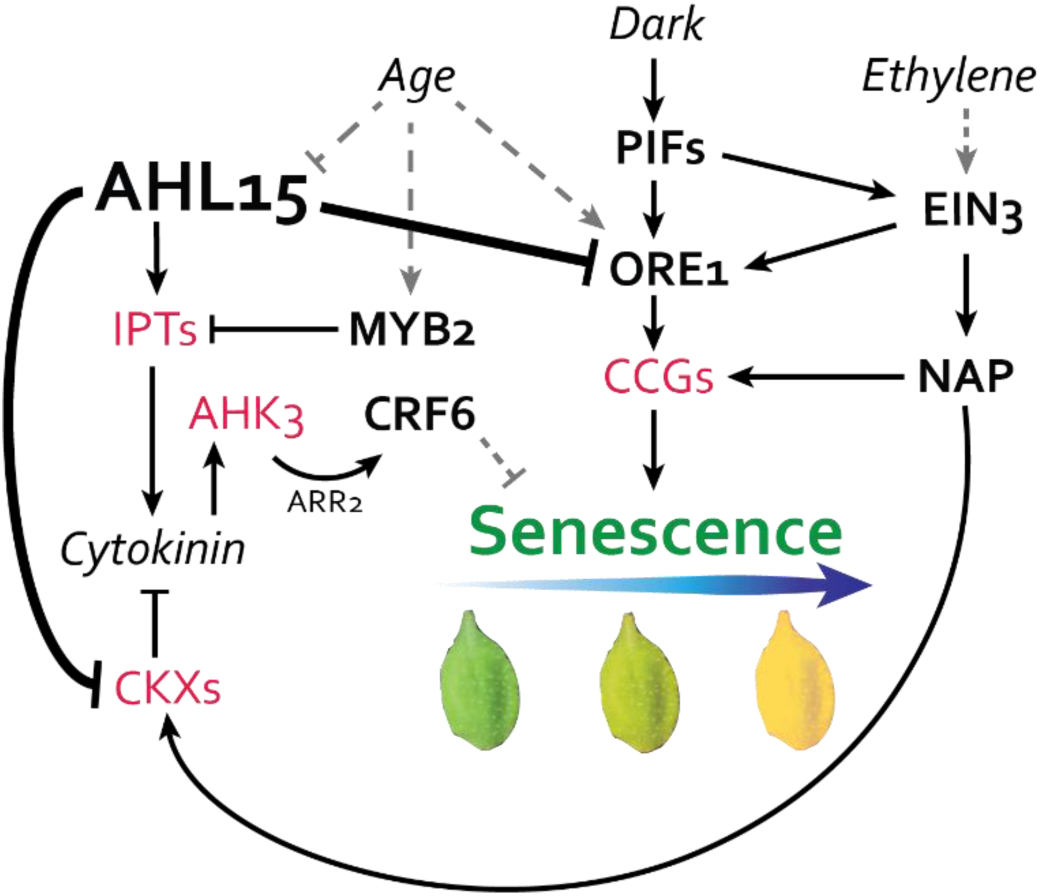
Model of AHL15-mediated repression of senescence. AHL15 represses ORESARA1 (ORE1), which is a key positive regulator of the senescence transcriptional network. ORE1 can be induced by dark incubation via the PHYTOCHROME INTERACTING FACTORS (PIFs), by ethylene via ETHYLENE INSENSITIVE 3 (EIN3), or by age. ORE1 activates several downstream targets among which chlorophyll catabolic genes (*CCG*s), ultimately leading to chlorophyll catabolism and causing the characteristic yellowing of senescent leaves. The NAC-LIKE, ACTIVATED BY AP3/PI (NAP) transcription factor acts in parallel to ORE1 and activates the expression of *CCG*s as well as the *CYTOKININ OXIDASE/DEHYDROGENASE* (*CKX*) genes, leading to the irreversible inactivation of cytokinins (CKs). CKs activate ARABIDOPSIS HISTIDINE KINASE 3 (AHK3), which represses leaf senescence via CYTOKININ RESPONSE FACTOR 6 (CRF6). CKs are produced by ISOPENTENYL TRANSFERASE (IPT) enzymes, which are transcriptionally repressed by MYB2 in ageing plants. AHL15 represses *CKX* expression and promotes the expression of *IPT7*. Lower CKX levels result in slower breakdown of CKs, resulting in higher CK levels and delayed senescence. TFs are indicated in bold letters, enzymes are indicated in red, and hormones and external abiotic factors are in italics.

## Discussion

AHL15 has previously been shown to delay several developmental processes, including vegetative phase change, flowering time, and axillary meristem maturation (Karami *et al*., 2020; Rahimi, Karami, Balazadeh, *et al*., 2022). Our data show that in addition to these processes, AHL15 delays leaf senescence by directly repressing the expression of *ORE1,* which encodes a key senescence regulator, and by limiting CK degradation through repression of *CKX* gene expression. These two processes are likely to act in parallel and together explain the strong stay-green phenotype observed in *AHL15* overexpressing plants (Figure 5). Moreover, our data suggest that AHL15 acts predominantly by repressing gene expression, as observed for *ORE1* and three *CKX* genes.

ORE1 acts as a key transcriptional regulator of senescence, and its expression is induced by several senescence-promoting environmental cues. Ethylene promotes the expression of *ORE1* and *NAP* via EIN3 (Li et al., 2013; Kim et al., 2014), ABA promotes *ORE1* expression via the TF ACTIVATING FACTOR1 (ATAF1) (Garapati *et al*., 2015), and dark incubation leads to enhanced *ORE1* transcription via the PIF TFs (Song *et al*., 2014). It was also shown that *ORE1* expression is post-transcriptionally silenced in young leaves by miR164 and that this silencing is relieved with progressing age (Kim *et al*., 2009; Li *et al*., 2013). Here, we have identified AHL15 as a novel repressor of *ORE1* expression, suggesting that AHL15 can antagonize the ethylene- and dark-induced senescence processes mediated by ORE1. In addition, *AHL15* is strongly expressed in juvenile plants and developing leaves and is gradually silenced over developmental age (Rahimi, Karami, Balazadeh, *et al*., 2022). It is therefore possible that AHL15 can act as an additional age-dependent repressor of *ORE1* expression in parallel to miR164 (Figure 5).

CK has long been recognized as an inhibitor of leaf senescence, yet its precise role and position within the senescence regulatory network remain relatively poorly understood. Application or overproduction of CK can block the onset of leaf senescence (Richmond *et al*., 1957; Gan and Amasino, 1995; Kim *et al*., 2006; Argueso and Kieber, 2024), and it has been shown that CK degradation is promoted during the senescence process by NAP-mediated upregulation of *CKX* expression (Hu *et al*., 2021; Li *et al*., 2023). MYB2 was shown to lower CK production in an age-dependent manner by repressing the expression of several *IPT* genes, which are the rate-limiting enzymes in the CK biosynthesis pathway (Guo and Gan, 2011). Our data show that AHL15 acts antagonistically to NAP in *CKX* transcriptional regulation and represses senescence via inhibition of CK degradation. In addition, we found that *IPT7* expression decreases in response to dark-induced senescence and that this process is delayed after activation of AHL15-GR by DEX treatment. Taken together, our results indicate that AHL15 acts as a positive regulator of CK in the senescence pathway, and thereby acts antagonistically to NAP and MYB2 in this process (Figure 5).

Our experiments show that the expression of the CK biosynthesis genes *LOG1* and *IPT7* is downregulated upon senescence induction by dark treatment. On the other hand, we observed an increased expression of *LOG8* after senescence induction, and that *LOG7* expression is stable throughout the senescence process. In addition, the levels of cZ in its base form and most of its derivatives were increased in senescent leaves. Together, these data show that CK biosynthesis is not completely repressed but partially maintained during the senescence developmental program. Our data also confirm the previously reported induction of *CKX* expression during senescence (Hu *et al*., 2021; Li *et al*., 2023). This indicates that CK levels are regulated both by biosynthesis and by degradation during the senescence process and suggests a role for some CK variants in the senescence process. Finally, our data show that inhibition of CK breakdown by direct silencing of *CKX* genes partially explains the delayed senescence phenotype of *AHL15* overexpressing plants. Previously, it was shown that expression of *LOG4*, *IPT3* and *IPT7* was decreased in *ahl15* knockout plants, resulting in lower CK levels and a decrease in radial xylem width (Rahimi, Karami, Lestari, *et al*., 2022). Here, we observed delayed downregulation of *IPT7* in dark-incubated leaves following DEX-induced nuclear localization of AHL15-GR, indicating that AHL15 targets are partially shared between stem and leaf tissue.

The leaf senescence phenotype of *ahl15* loss-of-function mutants is relatively mild (Figure S1), suggesting that either *AHL15* itself plays a minor role in leaf senescence, or that other *AHL* genes take over its role in the mutant plants. *AHL* genes have been shown to be highly functionally redundant, and developmental phenotypes are observed only in higher-order *ahl* mutants (Xiao *et al*., 2009; Zhao *et al*., 2013; Karami *et al*., 2020; Rahimi, Karami, Balazadeh, *et al*., 2022). In addition, *AHL15* is expressed primarily in young juvenile leaves as its expression in adult leaves is repressed by SPL TFs (Rahimi, Karami, Balazadeh, *et al*., 2022). In fact, with the *pAHL15:GUS* reporter, no expression is observed in fully expanded juvenile leaves or in adult leaves (Figure S2). It is therefore likely that higher-order *ahl* mutants, especially of the clade-A *AHL* genes that are expressed in adult leaves, show a stronger leaf senescence phenotype compared to the *ahl15* single mutant.

The functional redundancy among AHL proteins suggests that overexpression of other clade-A *AHLs* could have similar effects as AHL15 on the expression of *ORE1* and *CKX* genes and the overall CK levels in the plant, and that the repression of CK degradation is a general effect of high AHL dosage. Overexpression of at least one other clade-A *AHL, AHL27,* has been shown to inhibit senescence (Lim *et al*., 2007), indicating that, like flowering time, the senescence-inhibiting phenotype could be induced by multiple AHLs in this clade. It would therefore be interesting to see whether overexpression of other clade-A *AHLs* has a similar senescence phenotype as *AHL15* and *AHL27*, and whether this coincides with increased CK levels in the leaves. Arabidopsis contains 15 clade A *AHLs*, and the genomes of most crop species contain a similar or higher number of *AHL* genes (Zhao *et al*., 2014, 2020; Machaj and Grzebelus, 2021; W. M. Zhang *et al*., 2021; Wang *et al*., 2023). Breeding efforts to improve plant longevity and shelf life could focus on enhancing the expression of clade-A *AHLs,* which, due to their functional overlap and large number, represent promising targets for crop improvement.

## Materials and Methods

### Plant material and growth conditions

Seeds were sown on damp soil (90% turf soil with 10% sand) and stratified for three days at 4 °C before transfer to a climate chamber set at 21°C with a long day (16h photoperiod) and 65% relative humidity. Seedlings were transferred to individual pots one week after sowing and plants were watered weekly.

The *p35S:AHL15-GR* overexpression and *ahl15* loss-of-function lines have been described previously (Karami *et al*., 2020). *p35S:3xFLAG-AHL15* overexpression lines were generated by replacing the *AHL15-GR* construct in the *p35S:AHL15-GR* vector made by Karami et al. (2020) with a synthetic *3xFLAG-AHL15* construct, which was transformed to *Agrobacterium tumefaciens* strain AGL1 by electroporation (Dulk-Ras and Hooykaas, 1995). Transgenic *p35S:AHL15* Arabidopsis Col-0 lines were obtained by the floral dip method (Clough and Bent, 1998) and by subsequently selecting T1 and T2 plants on ½ MS medium supplemented with 15 mg/L phosphinotricin. Single locus transgenic lines, showing a 3:1 segregation ratio of resistant: susceptible plants in the T2 generation, were selected and T3 seeds from homozygous T2 plants were subsequently used for senescence experiments.

For dark-induced senescence assays, fifth leaves were detached from the plants at 23 days after the start of germination and floated on ca 5mL 5mM MES buffer (pH 5.6) in 5 cm ø Petri dishes, with a maximum of 10 leaves per dish. Petri dishes were sealed with parafilm, wrapped in aluminum foil, and stored in a cardboard box which was placed in the growth chamber to ensure a constant temperature of 20 °C. For the hormone treatments, a 10 mM dexamethasone (Thermo Fisher, Vilnius, Lithuania) stock solution dissolved in DMSO was diluted 1000-fold in 5 mM MES buffer to a final concentration of 10 μM. For mock treatment, DMSO was diluted 1000-fold in 5 mM MES buffer.

### Senescence measurements

For colorimetric measurements, the leaves were imaged on a dark background with a white reference card under uniform lighting with a Nikon DC3000 camera, with the following settings: ISO 160, exposure time 1/160 s. Image analysis was done in ImageJ with a custom plugin that measures Chl using RGB images (Luden *et al*., 2025). The analysis consisted of the following steps: white balancing of images based on a white reference sheet included in each picture; removal of background, conversion to an RGB image stack, automatic selection of individual leaves, and measurement of pixel intensity for each leaf in each of the red, green, and blue channels. The mean red, green, and blue values for each leaf were used to calculate the normalized red value (Rn; Red/(Red+Green+Blue)), which we previously showed correlates best with absolute Chl content (Luden *et al*., 2025).

After imaging for colorimetric measurements, two 5 mm ø leaf disks were taken from each leaf and stored at −80 °C. Acetone-based Chl extraction was done following the method described by Arnon (1949). Briefly, frozen leaf material was pulverized with a metal bead and dissolved in 200 μL 25 mM sodium phosphate buffer (pH 7) and 800 μL 80% acetone. Samples were then incubated at room temperature for 1 hour with gentle shaking and centrifuged for 10 minutes at 2500x*g*. 200 μL supernatant was then transferred to a clear-bottomed 96-well plate and the absorption (D) at 645 and 663 nm was measured with a TECAN Spark plate reader (Männedorf, Switzerland). The total Chl content in mg/L was then calculated with the following formula: Chl_Total_ = 20,2*D_645_ + 8,02*D_663_, as described by (Arnon, 1949), and then corrected for the total area of leaf tissue used for Chl extraction. Spectrophotometry results and colorimetric results from ImageJ were processed in R (R Core Team, 2023). Plots were generated using ggplot2 (Wickham, 2016) and ggpubr (Kassambara, 2023), and genotypes or treatments were compared with the Kruskal-Wallis test by using the package Agricolae (de Mendiburu, 2023).

### RNA isolation and RT-qPCR

After imaging for colorimetric measurements, leaves were cut in half, of which one half was used for RNA extraction, while the other half of each leaf was used for Chl extraction. For each RNA extraction, material from three plants was pooled and snap-frozen in liquid nitrogen, and tissue was pulverized with a metal bead using the Qiagen Tissuelyser. RNA was isolated with Trizol© reagent (Thermo Fisher Scientific, Vilnius, Lithuania) by following the standard protocol for Trizol-based RNA extraction, with an additional ethanol wash step at the RNA precipitation stage. Next, 1 μg of RNA was aliquoted to a new tube and treated with DNase I (Thermo Fisher Scientific, Vilnius, Lithuania) at 37 °C for 30 minutes and used for cDNA synthesis with the RevertAid First Strand cDNA Synthesis Kit (Thermo Fisher Scientific, Vilnius, Lithuania) according to the kit’s instructions. RT-qPCR was performed with TB green ex-Taq II (Takara Bio Europe, Saint-Germain-en-Laye, France) on the QuantStudio 5 Real-Time PCR machine (Applied Biosystems, Foster City, USA) in a reaction volume of 5 μL. Primers used for RT-qPCR (Supplementary Table 1) were synthesized by Sigma Aldrich (Haverhill, UK). Gene expression was normalized to *ACTIN2* expression by using the 2^-ΔΔCT method (Livak and Schmittgen, 2001), and groups were compared with the Kruskall-Wallis test in R as described above.

### Cytokinin measurements

Quantification of cytokinin metabolites was performed on biological replicates according to the method described by Svačinová et al. (2012), including modifications described by Antoniadi et al. (2015). Samples (10 mg fresh weight) were homogenized and extracted in 1 ml of modified Bieleski buffer (60% methanol, 10% formic acid and 30% water) together with a cocktail of stable isotope-labeled internal standards (0.2 pmol of CK bases, ribosides, N-glucosides, and 0.5 pmol of CK O-glucosides, nucleotides per sample added). The extracts were applied onto an Oasis MCX column (30 mg/1 ml, Waters) conditioned with 1 ml each of 100% methanol and water, equilibrated sequentially with 1ml of 50% (v/v) nitric acid, 1 ml of water, and 1 ml of 1M formic acid, and washed with 1 ml of 1M formic acid and 1 ml 100% methanol. Analytes were then eluted by two-step elution using 1 ml of 0.35M ammonium hydroxide aqueous solution and 2 ml of 0.35M NH4OH in 60% (v/v) methanol solution. The eluates were then evaporated to dryness in vacuo and stored at −20°C. Cytokinin levels were determined by an ultra-high performance liquid chromatography-electrospray tandem mass spectrometry (UHPLC-MS/MS) using stable isotope-labelled internal standards as a reference (Rittenberg and Foster, 1940). Separation was performed on an Acquity UPLC® i-Class System (Waters, Milford, MA, USA) equipped with an Acquity UPLC BEH Shield RP18 column (150×2.1 mm, 1.7 μm; Waters), and the effluent was introduced into the electrospray ion source of a triple quadrupole mass spectrometer Xevo™ TQ-S MS (Waters, Milford, MA, USA). Five independent biological replicates, each consisting of material from the fifth leaf of three plants, were performed, including two technical replicates of each.

## Data statement

Raw data and materials are available upon request to R.O.

## Acknowledgements

We thank Jan Vink and Altay Temel for plant care, and Ward de Winter, Ellora Basu, and Mariël Lavrijsen for technical support. We also thank Jelmer van Lieshout for feedback on the project and Marion Larue for help with pilot experiments. This work is part of the REJUVENATOR project with file number GSGT.2019.024 (to TL) of the Graduate School Green Top sectors research programme, which is financed by the Dutch Research Council (NWO).

## Author Contributions

The project was conceived by T. L. and R. O. with input from S. B.. All experiments were performed by T. L., except for the CK measurements, which were performed by P. A. and O. N.. Figures were prepared by T. L.. The manuscript was written by T. L. with input from R. O. and S. B., and the manuscript was read and approved by all authors.

## Supporting Information

**Figure S1:**
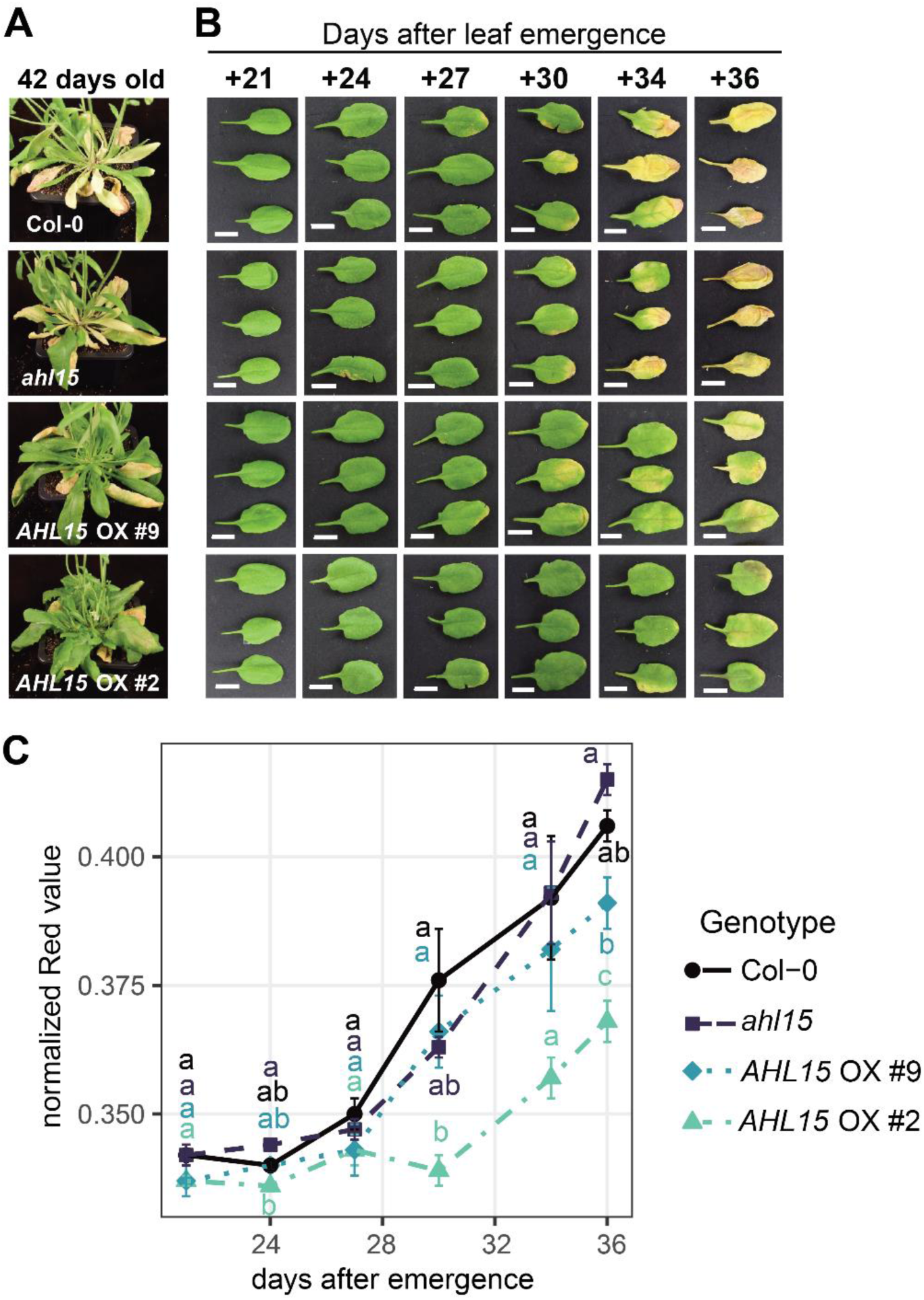
AHL15 delays age-dependent rosette leaf senescence. **A:** Images of 42-day-old wild-type (Col-0), *ahl15*, and *AHL15 OX* (*p35S:3xFLAG-AHL15*) plants grown in 5 cm-wide pots under long-day conditions. **B:** Progression of senescence in the fifth leaf of wild-type (Col-0), *ahl15*, and *AHL15 OX* (*p35S:3xFLAG-AHL15*) plants at different days after emergence. White bars represent 1 cm. **C:** Chlorophyll content was quantified based on the images in panel **B** as the normalized Red value (Rn), with four leaves per time point (only three leaves are shown in **B** to preserve space). Groups were compared at each time point by Kruskall-Wallis test, and letters indicate significance groups; groups with the same letter do not differ significantly from one another (p<0.05). Similar results were obtained in four independent experiments in different growth chambers.

**Figure S2:**
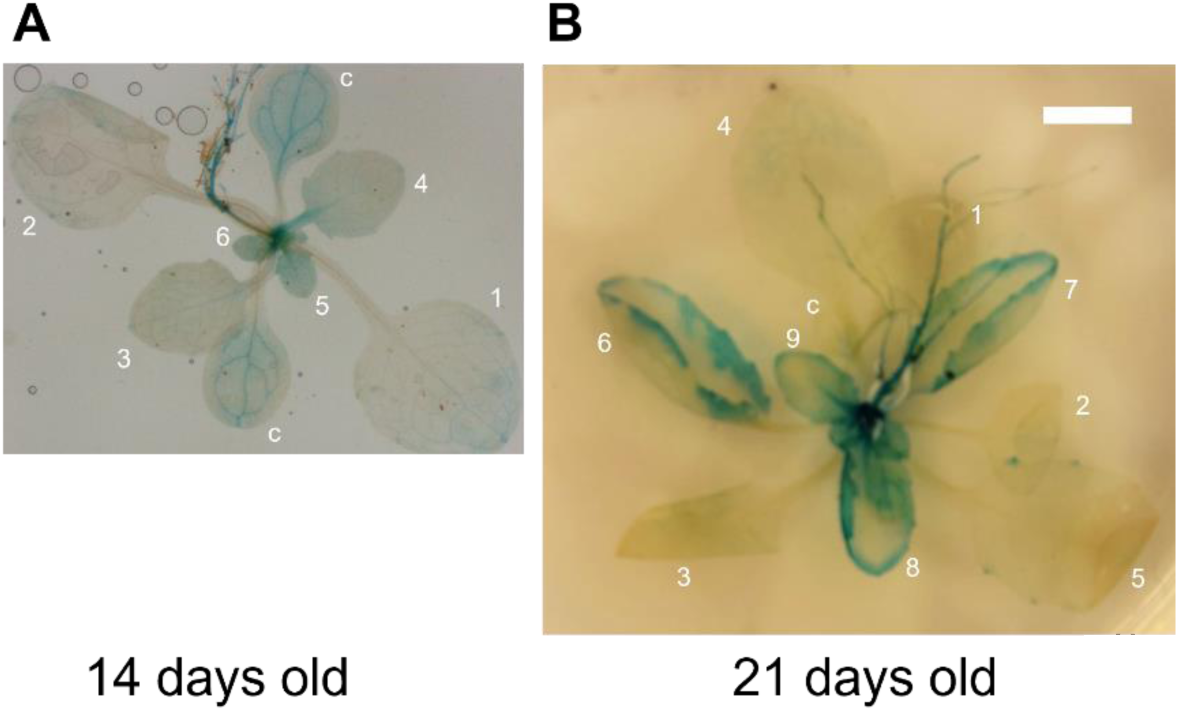
*AHL15* is expressed in young developing juvenile leaves. **A,B:** Images of 14-day-old (**A**) or 21-day-old (**B**) *pAHL15:GUS* GUS activity stained plants grown in long day (16h photoperiod) conditions. The leaf number (in order of appearance) is indicated with white numbers; cotyledons are marked with c. Size bar represents 5 mm.

**Figure S3.**
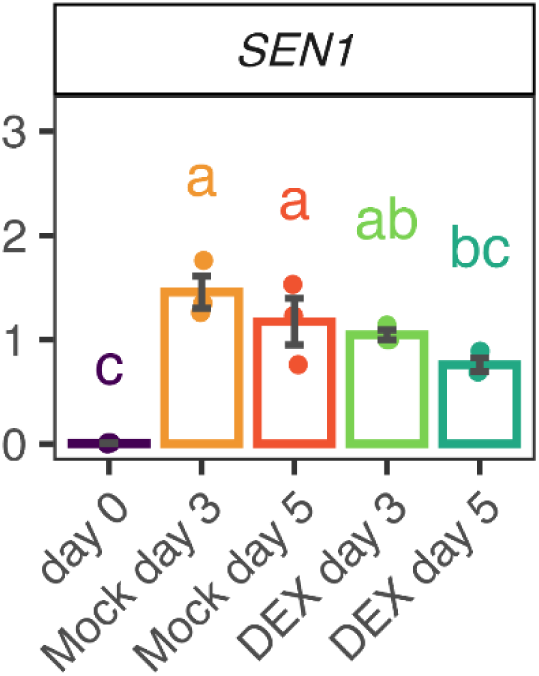
AHL15 only mildly reduces dark-induced *SEN1* expression in leaves. Fifth leaves were detached from 23-day-old *p35S:AHL15-GR* plants and incubated in the dark on 5 mM MES buffer with or without 10 µM DEX for the indicated number of days. Expression was determined by qRT-PCR in three biological replicates (n=3), each composed of leaf material of three plants, using *ACTIN2* as reference. The Kruskall-Wallis test was used to compare conditions, and significance groups are indicated by different letters (p < 0.05).

**Figure S4:**
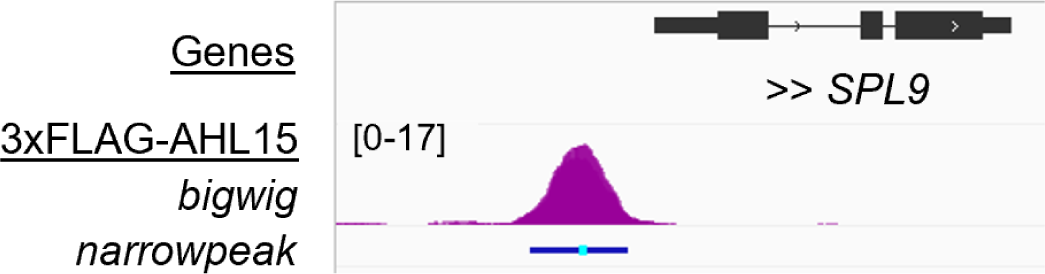
AHL15 binds upstream of the *SPL9* gene. ChIP seq performed on chromatin isolated from 31-day-old *p35S:3xFLAG-AHL15* plants using an anti-FLAG antibody for IP. The *SPL9* gene is shown in black (black blocks are exons, black lines are introns, transcription direction is indicated by a double arrow head), ChIP seq read alignments from the samples are shown in purple (bigwig track), and the location of significant peaks in blue, with a light blue bar indicating the peak summit (narrowpeak track) at approximately 500 bp upstream of the transcription start.

**Figure S5.**
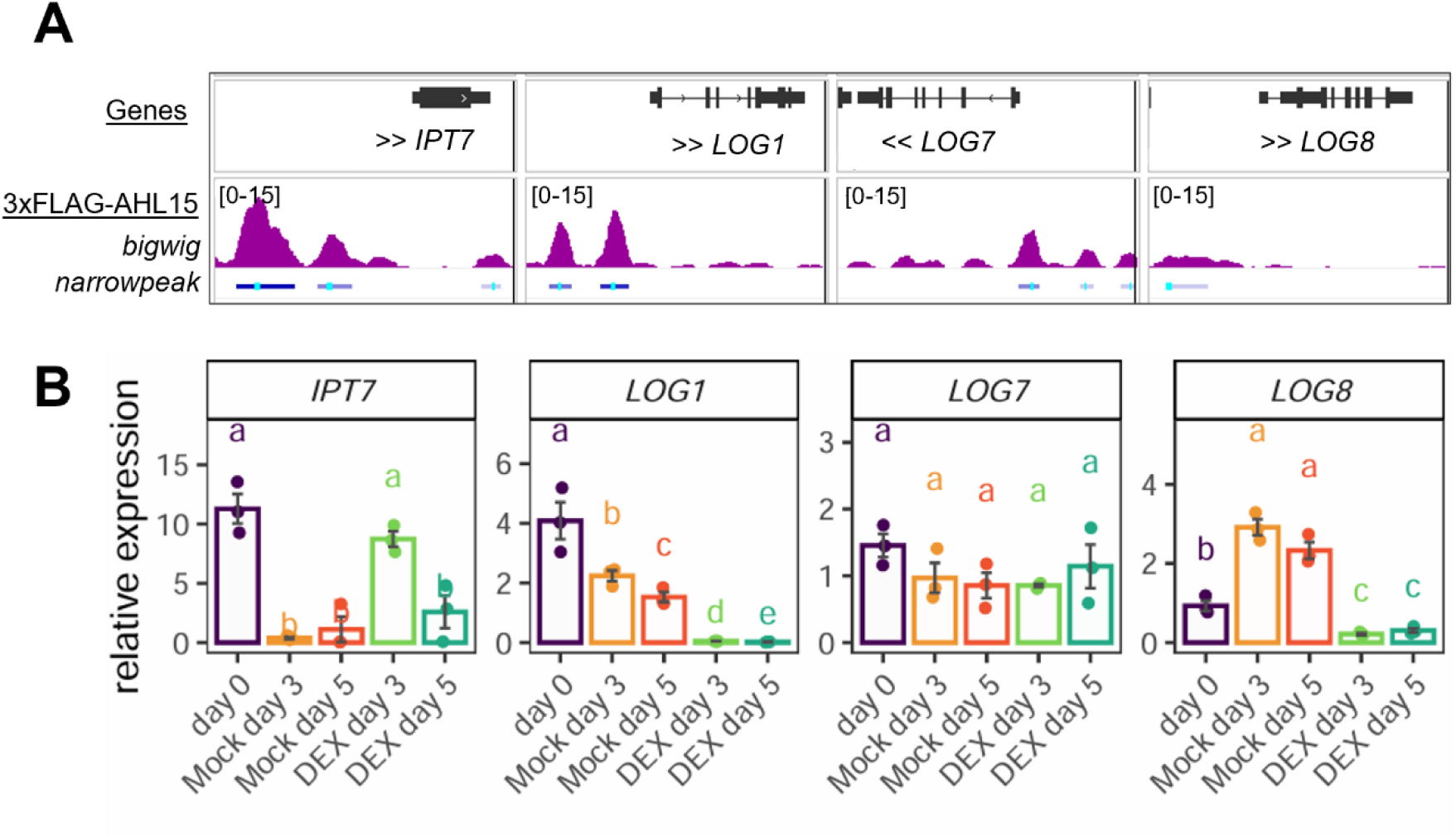
AHL15 binds to cytokinin biosynthesis genes to differentially regulate their expression. **A:** ChIP seq performed on chromatin isolated from 31-day-old *p35S:3xFLAG-AHL15* plants using an anti-FLAG antibody for IP. Genes are indicated black (black blocks are exons, black lines are introns), read alignments in purple (bigwig track), and the location of significant peaks in blue, with a light blue bar indicating the peak summit (narrowpeak track) at 1.4 kb and 3 kb upstream and in the 3’ UTR of *IPT7*, at 1 kb and 2.1 kb upstream of *LOG1*, directly upstream of *LOG7*, and at 1.6 kb upstream of *LOG8*. **B:** *IPT7, LOG1,LOG7,* and *LOG8* expression in the fifth leaf detached from 23-day old *p35S:AHL15-GR* plants and incubated in the dark on 5 mM MES buffer with or without 10 µM DEX for the indicated number of days. Expression was determined by qRT-PCR on three biological replicates (n=3), each composed of leaf material of three plants, using *ACTIN2* as reference. The Kruskall-Wallis test was used to compare conditions, and significance groups are indicated by different letters (p < 0.05).

**Figure S6.**
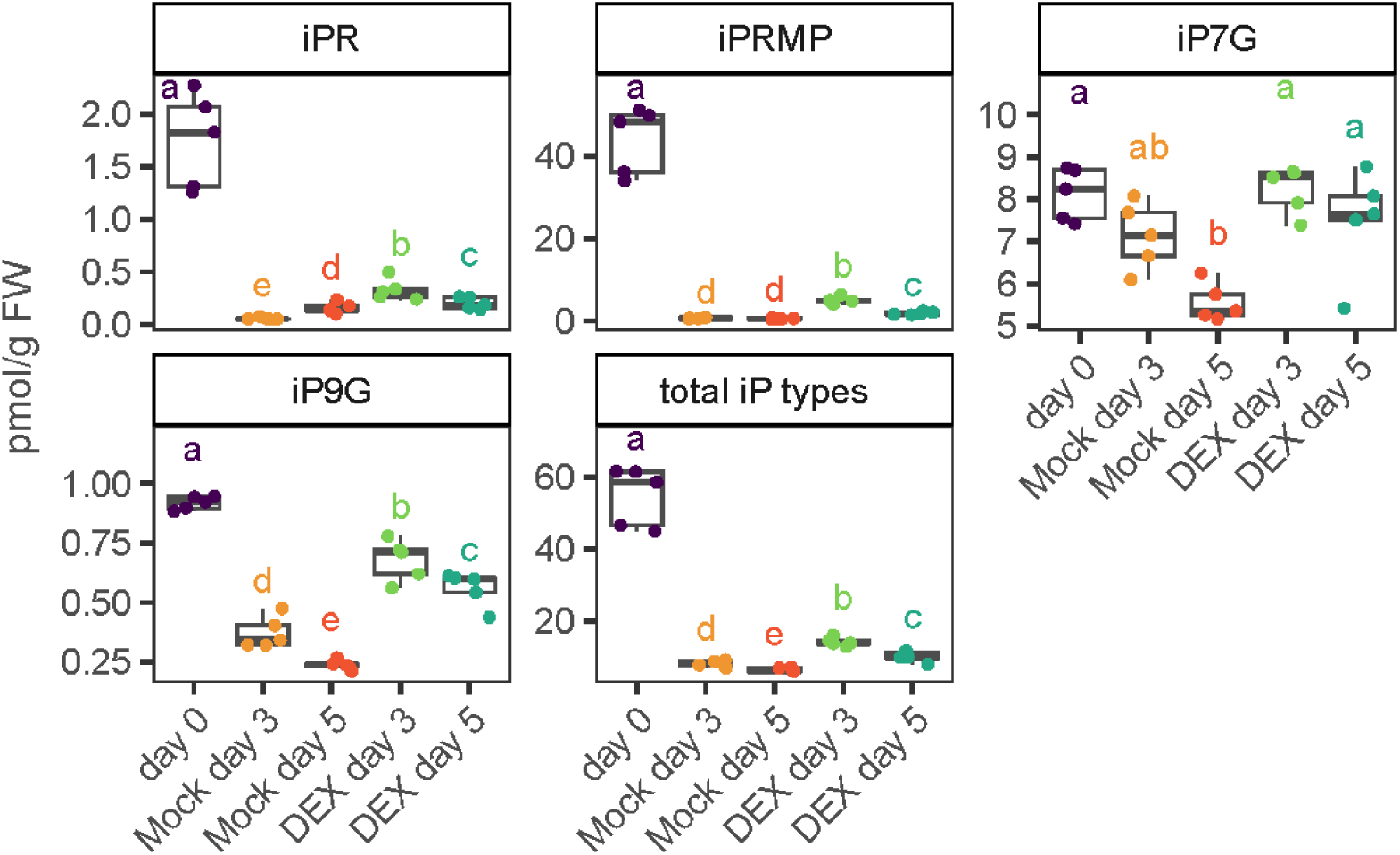
AHL15 delays the dark-induced decline in iP-type CK levels in leaves. The fifth leaf was collected from 23-day-old plants and floated on 5 mM MES with or without 10 µM DEX and incubated in the dark for the indicated number of days. The material of three leaves was pooled for each sample, and five samples were measured for each condition (n = 5). Groups were compared using the Kruskall-Wallis test, and letters indicate significance groups (p < 0.05). The iP base was not present at detectable levels.

**Figure S7.**
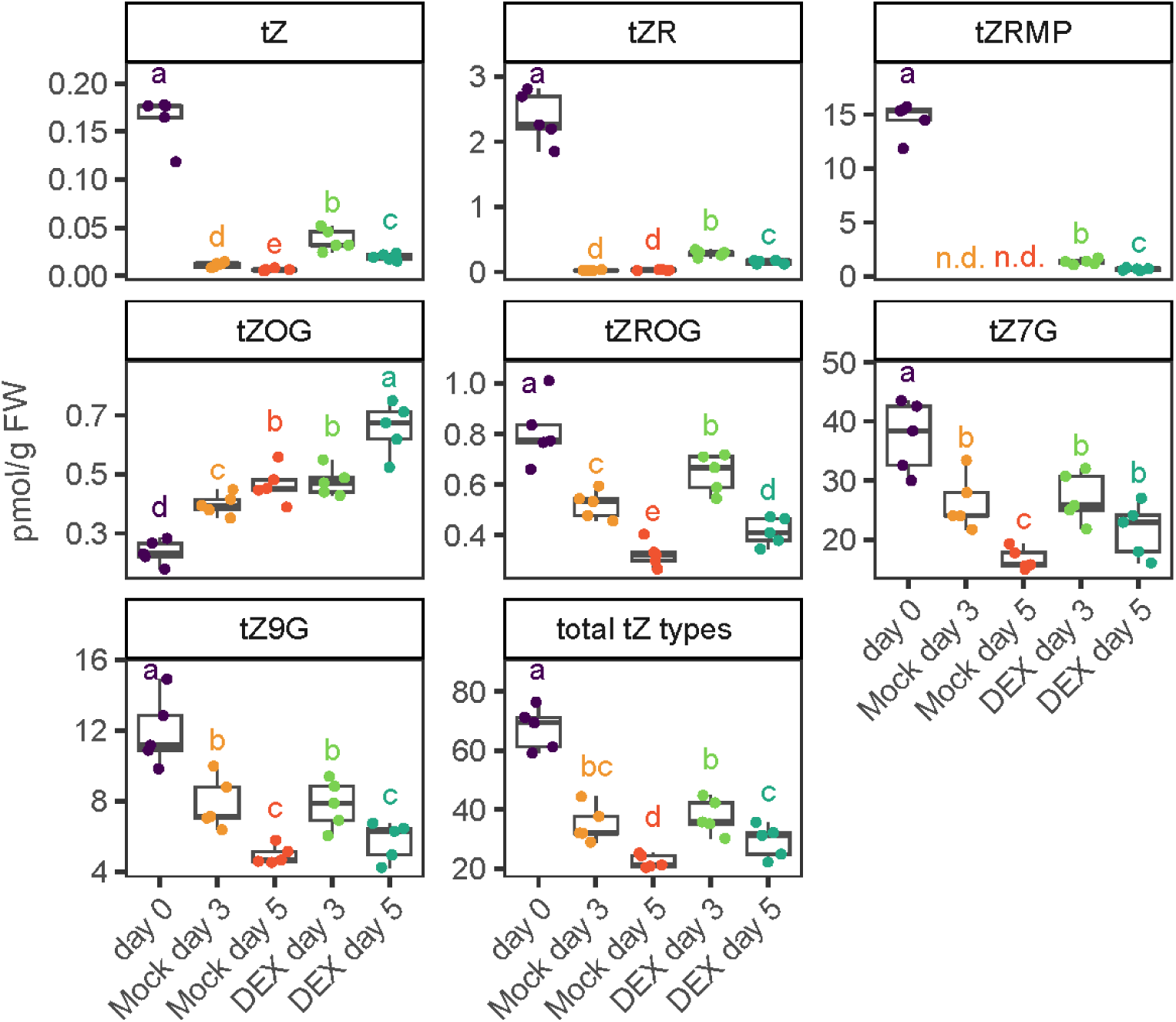
AHL15 delays the dark-induced decline in tZ- and tZR-type CK levels in leaves. The fifth leaf was collected from 23-day-old plants and floated on 5 mM MES with or without 10 µM DEX and stored in the dark for the indicated number of days. The material of three leaves was pooled for each sample, and five samples were measured for each condition (n = 5). Groups were compared using the Kruskall-Wallis test, and letters indicate significance groups (p < 0.05).

**Figure S8.**
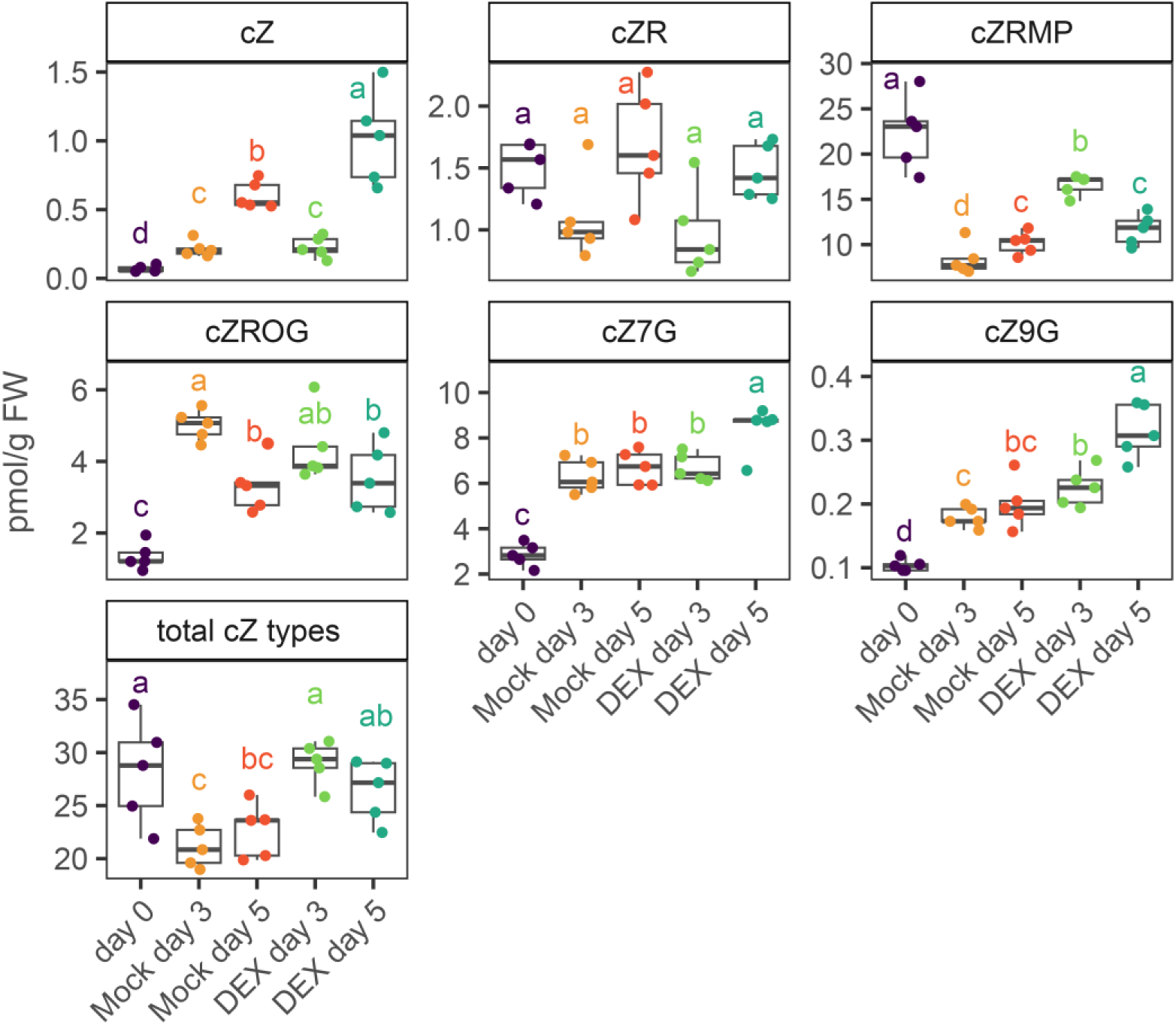
The effect of AHL15 on cZ-type CK levels in dark-incubated leaves. The fifth leaf was collected from 23-day-old plants and floated on 5 mM MES with or without 10 µM DEX and stored in the dark for the indicated number of days. The material of three leaves was pooled for each sample, and five samples were measured for each condition (n = 5). Groups were compared using the Kruskall-Wallis test, and letters indicate significance groups (p < 0.05).

**Figure S9.**
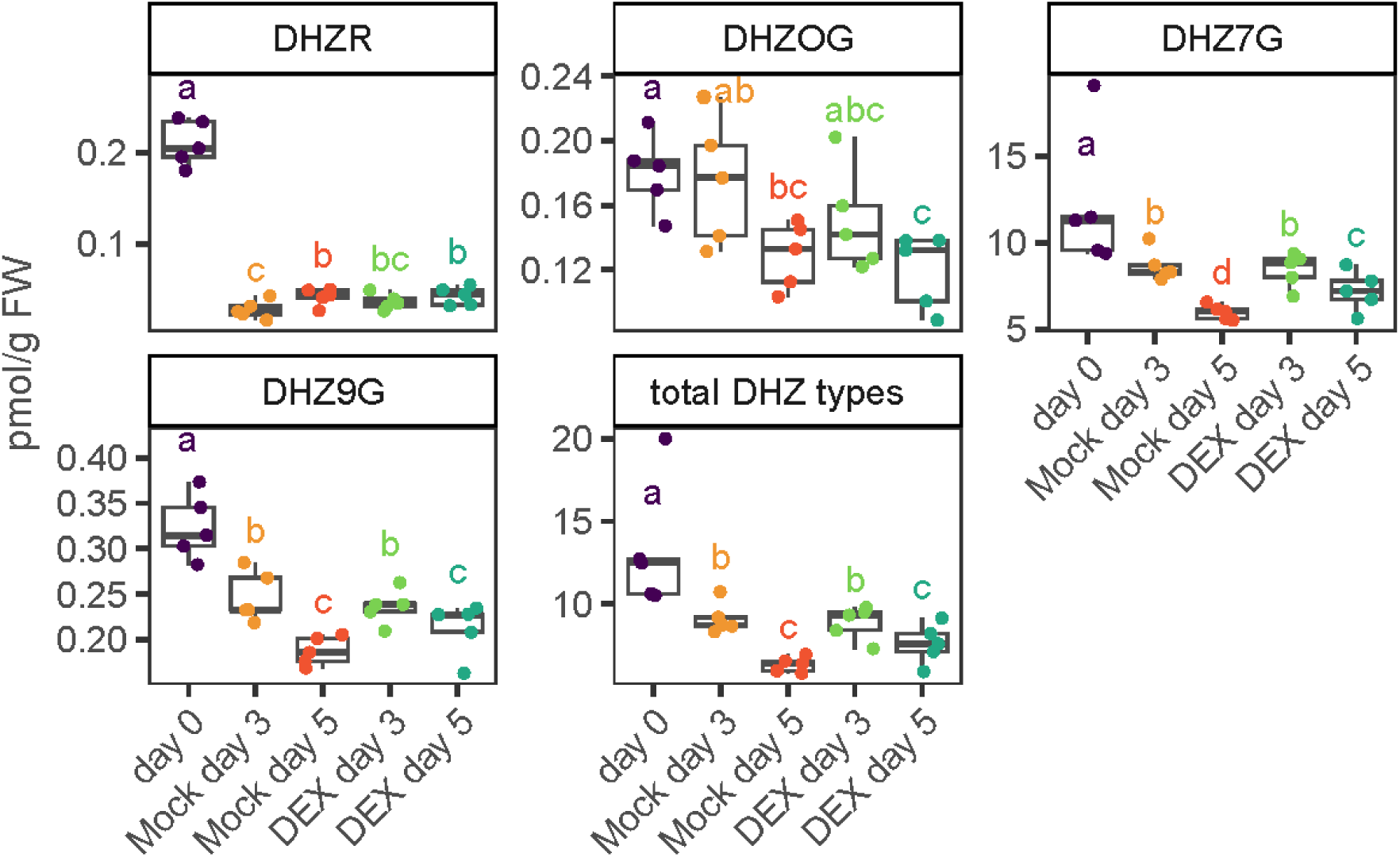
The effect of AHL15 on DHZ-type CK levels in dark-incubated leaves. The fifth leaf was collected from 23-day-old plants and floated on 5 mM MES with or without 10 µM DEX and stored in the dark for the indicated number of days. Material of three leaves was pooled for each sample, and five samples were measured for each condition (n = 5). Groups were compared using the Kruskall-Wallis test, and letters indicate significance groups (p < 0.05). DHZ base was not present at detectable levels in these samples.

**Figure S10.**
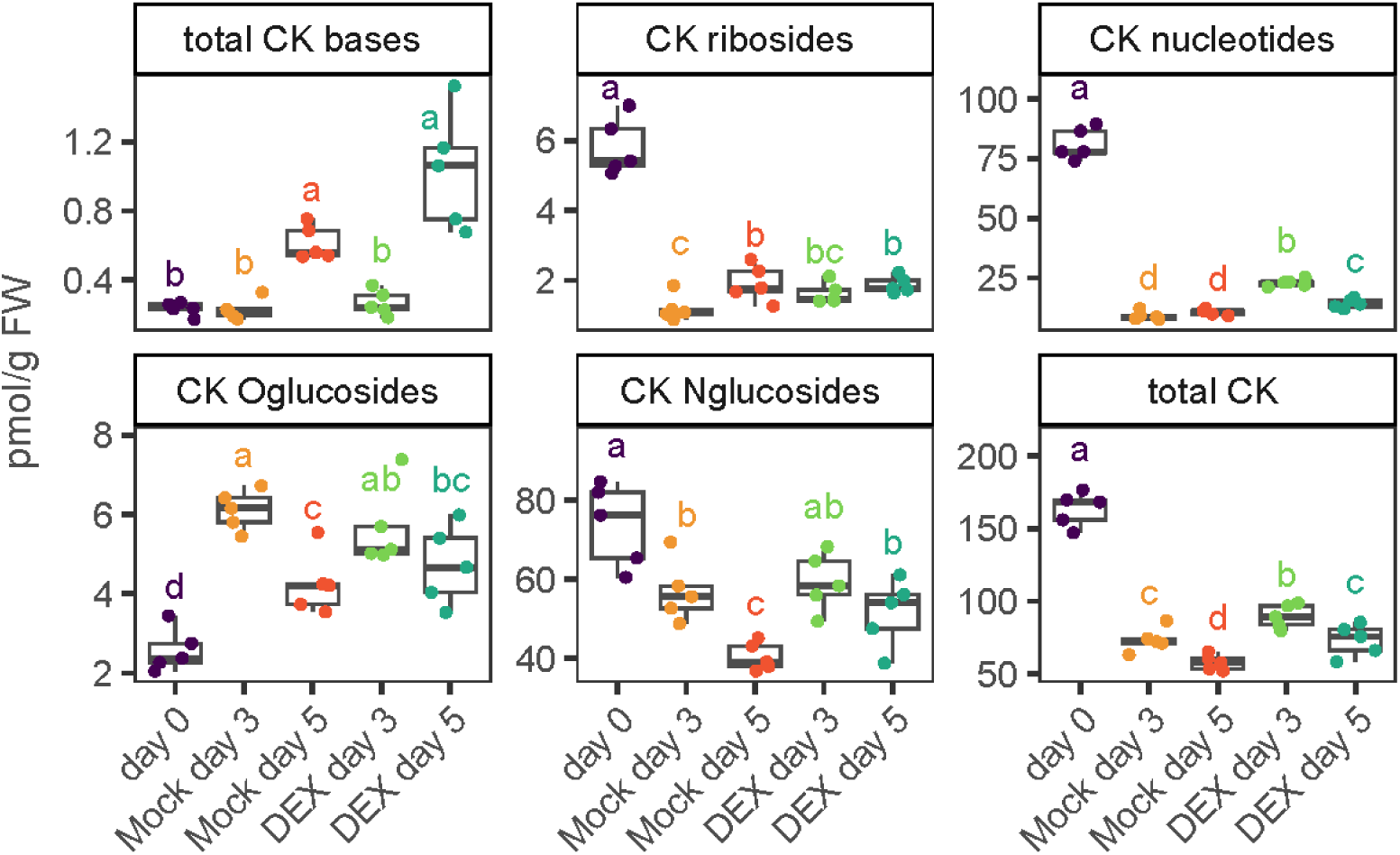
AHL15 delays the dark-induced decline in active CK levels in leaves. The fifth leaf was collected from 23-day-old plants and floated on 5 mM MES with or without 10 µM DEX and incubated in the dark for the indicated number of days. The material of three leaves was pooled for each sample, and five samples were measured for each condition (n = 5). Groups were compared using the Kruskall-Wallis test, and letters indicate significance groups (p < 0.05).

**Figure S11.**
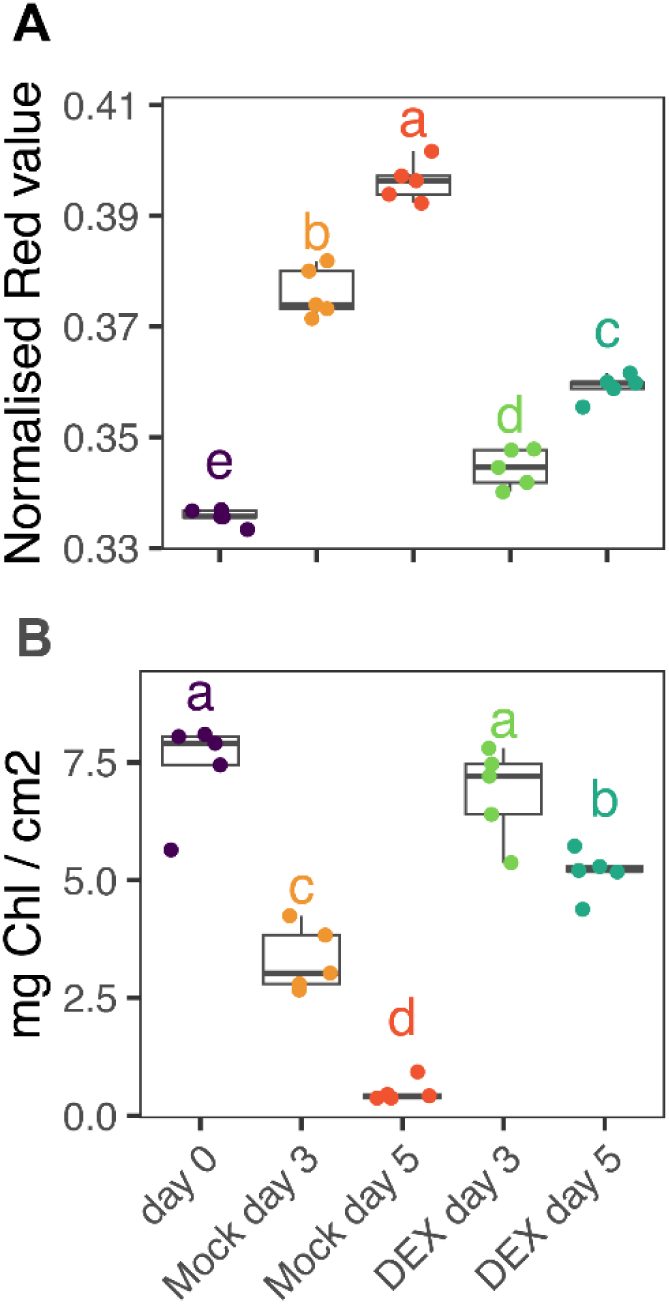
AHL15 preserves the chlorophyll content in leaves during dark incubation. **A:** Normalized Red values (Rn) based on RGB images of leaves just before collecting material for CK measurements. **B:** Total chlorophyll content determined by acetone-based chlorophyll extraction in leaf disks taken from the same samples as used for CK measurements. For each condition in **A** and **B**, five samples were measured (n = 5), each consisting of material from the fifth leaf harvested from three individual 23-day-old *p35S:AHL15-GR* plants. Groups were compared with the Kruskall-Wallis test, and letters indicate significance groups (p <0.05).

**Table S1:**
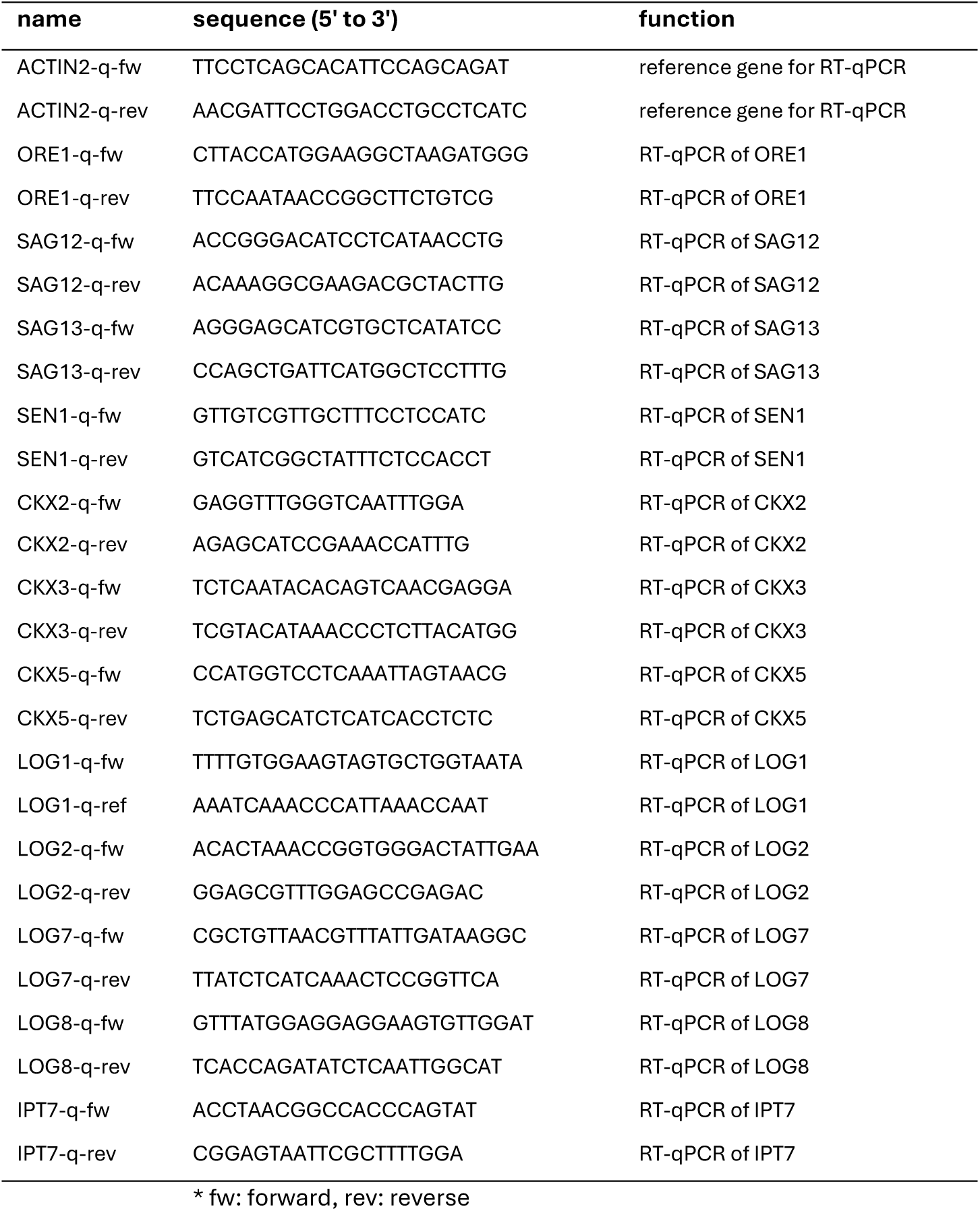
Primers used for RT-qPCR.

## Literature

Antoniadi, I. et al. (2015) ‘Cell-Type-Specific Cytokinin Distribution within the Arabidopsis Primary Root Apex’, Plant Cell, 27(7), pp. 1955–1967. Available at: 10.1105/TPC.15.00176.

Argueso, C.T. and Kieber, J.J. (2024) ‘Cytokinin: From autoclaved DNA to two-component signaling’, The Plant Cell, pp. 1–22. Available at: 10.1093/plcell/koad327.

Arnon, D.I. (1949) ‘Copper enzymes in isolated chloroplasts. Polyphenoloxidase in Beta Vulgaris’, Plant Physiology, 24(1), pp. 1–15. Available at: 10.1104/PP.24.1.1.

Balanzà, V. et al. (2018) ‘Genetic control of meristem arrest and life span in Arabidopsis by a FRUITFULL-APETALA2 pathway’, Nat. Commun., 9, p. 565. Available at: 10.1038/s41467-018-03067-5.

Balazadeh, S. et al. (2010) ‘A gene regulatory network controlled by the NAC transcription factor ANAC092/AtNAC2/ORE1 during salt-promoted senescence’, Plant J., 62(2), pp. 250–264. Available at: 10.1111/J.1365-313X.2010.04151.X.

Chung, B.-C. et al. (1997) ‘The promoter activity of sen 1, a senescence-associated gene of Arabidopsis, is repressed by sugars’, J. Plant Phys., 151, pp. 339–345. Available at: 10.1016/S0176-1617(97)80262-3.

Clough, S.J. and Bent, A.F. (1998) ‘Floral dip: a simplified method for Agrobacterium-mediated transformation of Arabidopsis thaliana’, Plant J., 16(6), pp. 735–743. Available at: 10.1046/J.1365-313X.1998.00343.X.

Dulk-Ras, A. den and Hooykaas, P.J.J. (1995) ‘Electroporation of Agrobacterium tumefaciens’, in Plant Cell Electroporation And Electrofusion Protocols. New Jersey: Humana Press, pp. 63–72. Available at: 10.1385/0-89603-328-7:63.

Gajdošová, S. et al. (2011) ‘Distribution, biological activities, metabolism, and the conceivable function of cis-zeatin-type cytokinins in plants’, J EXP BOT, 62(8), pp. 2827–2840. Available at: 10.1093/JXB/ERQ457.

Galuszka, P. et al. (2000) ‘Degradation of cytokinins by cytokinin oxidases in plants’, Plant Growth Regul., 32, pp. 315–327.

Gan, S. and Amasino, R.M. (1995) ‘Inhibition of leaf senescence by autoregulated production of cytokinin’, Science, 270(5244), pp. 1986–1988. Available at: 10.1126/SCIENCE.270.5244.1986.

Garapati, P. et al. (2015) ‘Transcription factor ATAF1 in Arabidopsis promotes senescence by direct regulation of key chloroplast maintenance and senescence transcriptional cascades’, Plant Physiol., 168(3), pp. 1122–1139. Available at: 10.1104/PP.15.00567.

Guo, Y. et al. (2021) ‘Leaf senescence: progression, regulation, and application’, Mol. Hortic., 1(1), pp. 1–25. Available at: 10.1186/S43897-021-00006-9.

Guo, Y. and Gan, S. (2006) ‘AtNAP, a NAC family transcription factor, has an important role in leaf senescence’, Plant J., 46(4), pp. 601–612. Available at: 10.1111/J.1365-313X.2006.02723.X.

Guo, Y. and Gan, S. (2011) ‘AtMYB2 regulates whole plant senescence by inhibiting cytokinin-mediated branching at late stages of development in arabidopsis’, Plant Physiol., 156(3), pp. 1612–1619. Available at: 10.1104/pp.111.177022.

Hallmark, H.T. and Rashotte, A.M. (2020) ‘Cytokinin isopentenyladenine and its glucoside isopentenyladenine-9G delay leaf senescence through activation of cytokinin-associated genes’, Plant Direct, 4(12), p. e00292. Available at: 10.1002/PLD3.292.

Hönig, M. et al. (2018) ‘Role of cytokinins in senescence, antioxidant defence and photosynthesis’, Int. J. Mol. Sci., 19(12). Available at: 10.3390/IJMS19124045.

Hu, Y. et al. (2021) ‘The leaf senescence-promoting transcription factor AtNAP activates its direct target gene CYTOKININ OXIDASE 3 to facilitate senescence processes by degrading cytokinins’, Mol. Hortic., 1(1), pp. 1–12. Available at: 10.1186/S43897-021-00017-6/FIGURES/7.

Jordi, W. et al. (2000) ‘Increased cytokinin levels in transgenic PSAG12–IPT tobacco plants have large direct and indirect effects on leaf senescence, photosynthesis and N partitioning’, Plant Cell Environ., 23(3), pp. 279–289. Available at: 10.1046/J.1365-3040.2000.00544.X.

Kakimoto, T. (2001) ‘Identification of Plant Cytokinin Biosynthetic Enzymes as Dimethylallyl Diphosphate:ATP/ADP Isopentenyltransferases’, Plant Cell Phys., 42(7), pp. 677–685. Available at: 10.1093/PCP/PCE112.

Karami, O. et al. (2020) ‘A suppressor of axillary meristem maturation promotes longevity in flowering plants’, Nat. Plants, 6(4), pp. 368–376. Available at: 10.1038/s41477-020-0637-z.

Kassambara, A. (2023) ggpubr: ‘ggplot2’ Based Publication Ready Plots. Available at: https://cran.r-project.org/package=ggpubr.

Kim, H.J. et al. (2006) ‘Cytokinin-mediated control of leaf longevity by AHK3 through phosphorylation of ARR2 in Arabidopsis’, PNAS, 103(3), pp. 814–819. Available at: 10.1073/PNAS.0505150103/SUPPL_FILE/05150TABLE1.PDF.

Kim, H.J. et al. (2014) ‘Gene regulatory cascade of senescence-associated NAC transcription factors activated by ETHYLENE-INSENSITIVE2-mediated leaf senescence signalling in Arabidopsis’, J EXP BOT, 65(14), pp. 4023–4036. Available at: 10.1093/JXB/ERU112.

Kim, J.H. et al. (2009) ‘Trifurcate feed-forward regulation of age-dependent cell death involving miR164 in Arabidopsis’, Science, 323(5917), pp. 1053–1057. Available at: 10.1126/SCIENCE.1166386/SUPPL_FILE/KIM-SOM.PDF.

Kuroha, T. et al. (2009) ‘Functional analyses of LONELY GUY cytokinin-activating enzymes reveal the importance of the direct activation pathway in Arabidopsis’, Plant Cell, 21, pp. 3152–3169. Available at: 10.1105/tpc.109.068676.

Li, F. et al. (2023) ‘A NAC transcriptional factor BrNAC029 is involved in cytokinin-delayed leaf senescence in postharvest Chinese flowering cabbage’, Food Chem., 404, p. 134657. Available at: 10.1016/j.foodchem.2022.134657.

Li, Z. et al. (2013) ‘ETHYLENE-INSENSITIVE3 Is a Senescence-Associated Gene That Accelerates Age-Dependent Leaf Senescence by Directly Repressing miR164 Transcription in Arabidopsis’, Plant Cell, 25(9), p. 3311. Available at: 10.1105/TPC.113.113340.

Lim, P.O. et al. (2007) ‘Overexpression of a chromatin architecture-controlling AT-hook protein extends leaf longevity and increases the post-harvest storage life of plants’, Plant J., 52(6), pp. 1140–1153. Available at: 10.1111/j.1365-313X.2007.03317.x.

Lim, P.O., Kim, H.J. and Nam, H.G. (2007) ‘Leaf Senescence’, Annu. Rev. Plant Biol., 58, pp. 115–136. Available at: 10.1146/annurev.arplant.57.032905.105316.

Livak, K.J. and Schmittgen, T.D. (2001) ‘Analysis of Relative Gene Expression Data Using Real-Time Quantitative PCR and the 2^-ΔΔCT Method’, Methods, 25(4), pp. 402–408. Available at: 10.1006/meth.2001.1262.

Lohman, K.N. et al. (1994) ‘Molecular analysis of natural leaf senescence in Arabidopsis thaliana’, Physiol. Plant., 92(2), pp. 322–328. Available at: 10.1111/J.1399-3054.1994.TB05343.X.

Luden, T. et al. (2025) ‘GreenLeafVI: A FIJI plugin for high-throughput analysis of leaf chlorophyll content’, bioRxiv, p. 2025.07.24.666635. Available at: 10.1101/2025.07.24.666635.

Luden, T., Chouaref, J. and Offringa, R. (2025) ‘AT-HOOK-MOTIF NUCLEAR LOCALIZED 15 extends plant longevity by binding at poorly accessible, epigenetic mark-depleted chromatin that surrounds transcribed regions’, bioRxiv, p. 2025.08.05.668696. Available at: 10.1101/2025.08.05.668696.

Machaj, G. and Grzebelus, D. (2021) ‘Characteristics of the at-hook motif containing nuclear localized (Ahl) genes in carrot provides insight into their role in plant growth and storage root development’, Genes, 12, p. 764. Available at: 10.3390/genes12050764.

McCabe, M.S. et al. (2001) ‘Effects of PSAG12-IPT gene expression on development and senescence in transgenic lettuce’, Plant Physiol., 127(2), pp. 505–516. Available at: 10.1104/PP.010244.

de Mendiburu, F. (2023) ‘agricolae: Statistical Procedures for Agricultural Research’. Available at: https://cran.r-project.org/package=agricolae.

Oh, S.A. et al. (1996) ‘A senescence-associated gene of Arabidopsis thaliana is distinctively regulated during natural and artificially induced leaf senescence’, Plant Mol. Biol., 30, pp. 739–754. Available at: 10.1007/BF00019008.

Pokorná, E. et al. (2021) ‘Cytokinin N-glucosides: Occurrence, Metabolism and Biological Activities in Plants’, Biomolecules, 11(1), p. 24. Available at: 10.3390/BIOM11010024.

Qiu, K. et al. (2015) ‘EIN3 and ORE1 Accelerate Degreening during Ethylene-Mediated Leaf Senescence by Directly Activating Chlorophyll Catabolic Genes in Arabidopsis’, PLoS Genet., 11(7), p. e1005399. Available at: 10.1371/journal.pgen.1005399.

R Core Team (2023) ‘R: A Language and Environment for Statistical Computing’. Vienna, Austria: R Foundation for Statistical Computing. Available at: https://www.r-project.org/.

Rahimi, A., Karami, O., Lestari, A.D., et al. (2022) ‘Control of cambium initiation and activity in Arabidopsis by the transcriptional regulator AHL15’, Curr. Biol., 32(8), pp. 1764–1775. Available at: 10.1016/J.CUB.2022.02.060.

Rahimi, A., Karami, O., Balazadeh, S., et al. (2022) ‘miR156 -independent repression of the ageing pathway by longevity-promoting AHL proteins in Arabidopsis’, New Phytol., 235, pp. 2424–2438. Available at: 10.1111/NPH.18292.

Richmond, A.E. et al. (1957) ‘Effect of Kinetin on Protein Content and Survival of Detached Xanthium Leaves’, Science, 125(3249), pp. 650–651. Available at: 10.1126/SCIENCE.125.3249.650.B.

Rittenberg, D. and Foster, G.L. (1940) ‘A NEW PROCEDURE FOR QUANTITATIVE ANALYSIS BY ISOTOPE DILUTION, WITH APPLICATION TO THE DETERMINATION OF AMINO ACIDS AND FATTY ACIDS’, J. Biol. Chem., 133(3), pp. 737–744. Available at: 10.1016/S0021-9258(18)73304-8.

Schippers, J.H.M. et al. (2015) ‘Living to die and dying to live: The survival strategy behind leaf senescence’, Plant Physiol., 169, pp. 914–930. Available at: 10.1104/pp.15.00498.

Schmülling, T. et al. (2003) ‘Structure and function of cytokinin oxidase/dehydrogenase genes of maize, rice, Arabidopsis and other species’, J Plant Res, 116, pp. 241–252. Available at: 10.1007/s10265-003-0096-4.

Song, Y. et al. (2014) ‘Age-Triggered and Dark-Induced Leaf Senescence Require the bHLH Transcription Factors PIF3, 4, and 5’, Mol. Plant, 7(12), pp. 1776–1787. Available at: 10.1093/MP/SSU109.

Street, I.H. et al. (2008) ‘The AT-hook-containing proteins SOB3/AHL29 and ESC/AHL27 are negative modulators of hypocotyl growth in Arabidopsis’, Plant Journal, 54(1), pp. 1–14. Available at: 10.1111/j.1365-313X.2007.03393.x.

Svačinová, J. et al. (2012) ‘A new approach for cytokinin isolation from Arabidopsis tissues using miniaturized purification: pipette tip solid-phase extraction’. Available at: 10.1186/1746-4811-8-17.

Takei, K., Sakakibara, H. and Sugiyama, T. (2001) ‘Identification of genes encoding adenylateisopentenyltransferase, a cytokinin biosynthesis enzyme, in Arabidopsis thaliana’, J. Biol. Chem., 276(28), pp. 26405–26410. Available at: 10.1074/JBC.M102130200.

Thomas, H. (2013) ‘Senescence, ageing and death of the whole plant’, New Phytol., 197, pp. 696–711. Available at: 10.1111/nph.12047.

Tokunaga, H. et al. (2012) ‘Arabidopsis lonely guy (LOG) multiple mutants reveal a central role of the LOG-dependent pathway in cytokinin activation’, Plant J., 69(2), pp. 355–365. Available at: 10.1111/J.1365-313X.2011.04795.X.

Wang, L. et al. (2023) ‘Identification of tomato AHL gene families and functional analysis their roles in fruit development and abiotic stress response’, Plant Physiology and Biochemistry, 202(May), p. 107931. Available at: 10.1016/j.plaphy.2023.107931.

Wickham, H. (2016) ggplot2: Elegant Graphics for Data Analysis. Springer-Verlag New York. Available at: https://ggplot2.tidyverse.org (Accessed: 4 July 2024).

Woo, H.R. et al. (2019) ‘Leaf Senescence: Systems and Dynamics Aspects’, Annu. Rev. Plant Biol., 70, pp. 347–376. Available at: 10.1146/annurev-arplant-050718-095859.

Xiao, C. et al. (2009) ‘Over-expression of an AT-hook gene, AHL22, delays flowering and inhibits the elongation of the hypocotyl in Arabidopsis thaliana’, Plant Mol. Biol., 71(1–2), pp. 39–50. Available at: 10.1007/S11103-009-9507-9/FIGURES/6.

Yang, J., Worley, E. and Udvardi, M. (2015) ‘A NAP-AAO3 Regulatory Module Promotes Chlorophyll Degradation via ABA Biosynthesis in Arabidopsis Leaves’, Plant Cell, 26(12), pp. 4862–4874. Available at: 10.1105/TPC.114.133769.

Yang, T. et al. (2022) ‘The S40 family members delay leaf senescence by promoting cytokinin synthesis’, Plant Physiol. Biochem., 191, pp. 99–109. Available at: 10.1016/J.PLAPHY.2022.09.017.

Yun, J. et al. (2012) ‘The AT-hook motif-containing protein AHL22 regulates flowering initiation by modifying FLOWERING LOCUS T chromatin in Arabidopsis’, J. Biol. Chem., 287(19), pp. 15307–15316. Available at: 10.1074/jbc.M111.318477.

Zhang, W. et al. (2021) ‘Cytokinin oxidase/dehydrogenase OsCKX11 coordinates source and sink relationship in rice by simultaneous regulation of leaf senescence and grain number’, Plant Biotechnol. J., 19(2), pp. 335–350. Available at: 10.1111/PBI.13467.

Zhang, W.M. et al. (2021) ‘Insights into the molecular evolution of AT-Hook Motif nuclear Localization genes in Brassica napus’, Front. Plant Sci., 12. Available at: 10.3389/fpls.2021.714305.

Zhao, J. et al. (2013) ‘Arabidopsis thaliana AHL family modulates hypocotyl growth redundantly by interacting with each other via the PPC/DUF296 domain’, PNAS, 110(48), pp. E4688–E4697. Available at: 10.1073/pnas.1219277110.

Zhao, J. et al. (2014) ‘Insights into the evolution and diversification of the AT-hook Motif Nuclear Localized gene family in land plants’, BMC Plant Biol., 14(1), pp. 1–19. Available at: 10.1186/s12870-014-0266-7.

Zhao, L. et al. (2020) ‘Genome-wide identification and analyses of the AHL gene family in cotton (Gossypium)’, BMC Genomics, 21(1). Available at: 10.1186/s12864-019-6406-6.

Zhou, Y. et al. (2022) ‘Overexpression of AHL9 accelerates leaf senescence in Arabidopsis thaliana’, BMC Plant Biol., 22(1), pp. 1–12. Available at: 10.1186/S12870-022-03622-9.

Zwack, P.J. et al. (2013) ‘Cytokinin Response Factor 6 negatively tegulates leaf senescence and is induced in response to cytokinin and numerous abiotic stresses’, Plant Cell Phys., 54(6), pp. 971–981. Available at: 10.1093/PCP/PCT049.

